# Transcriptomic profiling of sex-specific olfactory neurons reveals subset-specific receptor expression in *Caenorhabditis elegans*

**DOI:** 10.1101/2021.10.26.465928

**Authors:** Douglas K. Reilly, Erich M. Schwarz, Caroline S. Muirhead, Annalise N. Robidoux, Anusha Narayan, Meenakshi K. Doma, Paul W. Sternberg, Jagan Srinivasan

**Author notes:** Corresponding author: Jagan Srinivasan. These authors contributed equally to this work (Douglas K. Reilly, Erich M. Schwarz).

## Abstract

**Summary:** The nematode *Caenorhabditis elegans* utilizes chemosensation to navigate an ever-changing environment for its survival. A class of secreted small-molecule pheromones, termed ascarosides, play an important role in olfactory perception by affecting biological functions ranging from development to behavior. The ascaroside ascr#8 mediates sex-specific behaviors, driving avoidance in hermaphrodites and attraction in males. Males sense ascr#8 via the ciliated male-specific cephalic sensory (CEM) neurons, which exhibit radial symmetry along dorsal-ventral and left-right axes. Calcium imaging studies suggest a complex neural coding mechanism that translates stochastic physiological responses in these neurons to reliable behavioral outputs. To test the hypothesis that neurophysiological complexity arises from differential expression of genes, we performed cell-specific transcriptomic profiling; this revealed between 18 and 62 genes with at least two-fold higher expression in a specific CEM neuron type versus both other CEM neurons and adult males. These included two G protein coupled receptor (GPCR) genes, *srw-97* and *dmsr-12*, that were specifically expressed in non-overlapping subsets of CEM neurons and whose expression was confirmed by GFP reporter analysis. Single CRISPR-Cas9 knockouts of either *srw-97* or *dmsr-12* resulted in partial defects, while a double knockout of both *srw-97* and *dmsr-12* completely abolished the attractive response to ascr#8. Together, our results suggest that the evolutionarily distinct GPCRs SRW-97 and DMSR-12 act non-redundantly in discrete olfactory neurons to facilitate male-specific sensation of ascr#8.

## Introduction

The ability of an organism to find a mate is critical to the survival of a species. Many species utilize small molecule pheromones to signal mate location (Pungaliya *et al*. 2009; Narayan *et al*. 2016), sexual maturity (Aprison AND Ruvinsky 2015; Aprison AND Ruvinsky 2017), and receptivity (Houck *et al*. 2007; Jang *et al*. 2017), which are sensed and processed via the nervous system, driving behavioral and developmental responses.

The nematode *Caenorhabditis elegans* communicates with conspecifics almost exclusively using pheromones called ascarosides (Ludewig *et al*. 2019; Mcgrath and Ruvinsky 2019), a structurally conserved, modular class of small molecule pheromones (Vonreuss and Schroeder 2015; Zhang *et al*. 2017) composed of a core ascarylose sugar and a fatty-acid derived side chain (Butcher *et al*. 2007; Von Reuss *et al*. 2012; Ludewig *et al*. 2019). Ascarosides signal a host of environmental and developmental information, including the sexual maturity and location of potential mates (Narayan *et al*. 2016; Aprison and Ruvinsky 2017), and this information is often conserved across nematodes (Choe *et al*. 2012; Ragsdale *et al*. 2013; Dong *et al*. 2016; Dong *et al*. 2018; Reilly *et al*. 2019).

While most male-specific neurons are located in the tail, the major contributor to male-specific chemosensory-driven behaviors are the CEM neurons, located in the amphid. Previous studies have shown these four radially symmetric neurons to be involved in sensing ascaroside #3 and #8 (ascr#3 and ascr#8), two pheromones that serve to attract males to mates (Pungaliya *et al*. 2009; Narayan *et al*. 2016; Reilly *et al*. 2017). Ascr#3 also plays a role in dauer formation, functioning alongside ascr#2 and ascr#4 as the “dauer pheromone” (Butcher *et al*. 2008). Ascr#8 is unique in that it contains a *p*-aminobenzoic acid moiety on its terminus (Pungaliya *et al*. 2009; Artyukhin *et al*. 2018). In animals lacking CEM the attractive response to ascr#8 is abolished, while further ablation of the ASK is required for complete loss of the ascr#3 behavioral response (Narayan *et al*. 2016). Ablating three of the four CEM neurons, while leaving one CEM intact, revealed that the four CEM neurons cooperate to drive a tuned attractive response to an intermediate (1 μM) concentration of both ascr#3 and ascr#8 (Narayan *et al*. 2016).

Calcium imaging and electrophysiological experiments demonstrated that CEM neurons show variable responses to stimuli not only between different animals, but within a single animal (Narayan *et al*. 2016; Reilly *et al*. 2017). While neurons such as AWC exhibit variable calcium responses between different animals, such responses are consistent between the left and right AWC neurons of a single animal (Cochella *et al*. 2014). In contrast, the responses of individual CEM neurons are stochastic within single animals, and yet four CEM neurons within one animal consistently generate proper behavioral responses (Narayan *et al*. 2016). To understand this pattern of stochastic neuronal activity yielding consistent behavioral outputs, it is imperative to uncover genes encoding components of the CEM response.

A recent study in *C. elegans* hermaphrodites generated transcriptomic landscapes of 118 neuronal classes from 302 neurons in order to link functional and anatomical properties of individual neurons with their molecular identities (Taylor *et al*. 2021). Discrete neuronal classes were successfully identified via their combinations of expressed neuropeptides and neuropeptide receptor genes. However, a similar feat has yet to be performed on male *C. elegans*, and the transcriptomic profiles of individual male neurons remain enigmatic.

In a more focused transcriptomic approach, gene expression profiles of extracellular vesicle-releasing neurons (EVNs) were identified (Wang *et al*. 2015; Kaletsky *et al*. 2016). While CEM neurons are a subset of EVN neurons, and CEM transcriptomic data are embedded in these data, there were thousands of EVNs in each replicate, diluting CEM-specific expression values.

To begin understanding how CEM neurons achieve stochastic yet reliable physiological responses to ascr#8, we performed single-cell RNA-seq (Schwarz *et al*. 2012) on CEM neurons, and uncovered a small number of highly enriched genes encoding G protein-coupled receptors (GPCRs). Given that all ascaroside receptors identified to date have been GPCRs (Kim *et al*. 2009; Mcgrath *et al*. 2011; Park *et al*. 2012; Greene *et al*. 2016a; Greene *et al*. 2016b; Chute *et al*. 2019), we tested whether any of these enriched genes contribute to male *C. elegans* sensation of ascr#8. We identified two distantly related GPCR genes that contribute to the ascr#8 behavioral response, *srw-97* and *dmsr-12*. Loss of each receptor resulted in partially deficient behavioral responses to ascr#8. Following generation of a double *srw-97;dmsr-12* mutant, we found that loss of both receptors results in a complete loss of behavioral response to ascr#8. Phylogenetic analysis further indicates that both receptors are homologous to closely related receptors present throughout the *Caenorhabditis* genus, and robust ascr#8 responses may be a trait recently evolved in *C. elegans*.

## Results

### The transcriptomic landscape of CEM neurons is variable

Individual CEM neurons were isolated from *C. elegans* expressing an integrated GFP labeling male-specific EVNs (p*pkd-2*::GFP [**Figure 2A**]), as previously described (Goodman *et al*. 1998; Narayan *et al*. 2011; Narayan *et al*. 2016). Cells were separated by anatomical identity (i.e., CEM dorsal left [DL], dorsal right [DR], ventral left [VL], and ventral right [VR]); single-cell CEM cDNA libraries were constructed and used for RNA-seq (Schwarz *et al*. 2012).

Enriched genes in each CEM neuron were identified by comparing RNA-seq profiles of distinct CEM types to one another and to whole adult *C. elegans* males (**Table S1, S5**). We observed 267 genes with at least two-fold higher expression in CEM neurons than in adult males. Of these, we observed 164 genes that were specific to individual CEM types, defining specificity as being at least two-fold higher expression in one type than in any other type: 62 genes specific to CEM DL, 40 to CEM DR, 18 to CEM VL, and 44 to CEM VR. Uniquely mapped reads ranged from 5.2 to 9.2 million per CEM type, matching the trend for alignment rates of each CEM neuron, which ranged from 18.0% to 47.2%, (**Table S2**) with an average of 1,426 genes showing robust RNA expression in each neuron (**Table S3**).

Five GPCR-encoding genes expressed in CEMs (*seb-3, srr-7, srw-97, dmsr-12, srd-32*) were selected for study; of these, *srw-97* and *dmsr-12* showed specific expression in CEM VR and CEM DL. Four of these GPCR genes were uncharacterized; one (*seb-3*) has previously been shown to play roles in locomotion, stress response, and ethanol tolerance (Jee *et al*. 2013). *dmsr-12* is related to *daf-37* (Robertson and Thomas 2006), a previously identified ascaroside receptor gene (Park *et al*. 2012), although it is more closely related to the neuropeptide receptor gene *dmsr-1*, and more distantly to *srw-97* (Robertson and Thomas 2006). *srd-32* belongs to a divergent branch of the SRD phylogeny (Robertson and Thomas 2006). *srr-7* belongs to the one of the smallest families of *C. elegans* chemoreceptor genes (Robertson and Thomas 2006).

*seb-3* and *srw-97* exhibited similar enrichment profiles across the CEM neurons, with 3.0- and 3.3-fold higher expression in CEM VR than in any other CEM neuron (**Table 1; Figure 1D; Table S5**). *dmsr-12* showed 25-fold higher expression in CEM DL versus any other CEM neuron (**Table 1; Figure 1A; Table S5**), while *srd-32* was only 1.8-fold higher in CEM DL (**Table 1; Figure 1A; Table S5**). *srr-7* showed 4.2-fold higher expression in CEM VL versus any other CEM neuron (**Table 1; Figure 1C; Table S5**), which correlated with previous transcriptomic analyses that found *srr-7* to be enriched in *C. elegans* EVNs (Wang *et al*. 2015).

**Table 1.** Expression levels for genes of interest in CEM neurons. For each gene, observed RNA-seq expression levels in each CEM neuron type are given in transcripts per million (TPM); CEM types with the highest expression are in boldface for each gene. Two measures of gene enrichment and specificity are also shown: first, the ratio of a gene’s expression for the highest-expressing CEM neuron type divided by that gene’s expression in the highest-expressing of the other three CEM neuron types (‘CEM/others’; ‘CEM_any/CEM_non_any.max’ in **Table S5**); and, the ratio of that gene’s highest-expressing CEM neuron type divided by that gene’s expression in the highest-expressing of three replicates of adult *C. elegans* males (‘CEM/males’; ‘CEM_all.max_TPM/male.max_TPM’ in **Table S5**). The first ratio gives the skew for a gene’s expression towards a single CEM type; the second ratio gives the degree to which that gene’s expression cannot be explained merely by its being a male-specific gene, perhaps expressed throughout the male body. Genes listed here are of interest either because we tested them for transgenic GFP expression or biological function, because they have previously known CEM-specificity, or because they are evolutionary related to *srw-97* and *dmsr-12*.

| Gene | CEM DL | CEM DR | CEM VL | CEM VR | CEM/others | CEM/males |
| --- | --- | --- | --- | --- | --- | --- |
| *srw-97* | 0.02 | 0.03 | 0.02 | **0.10** | 3.33 | 2.50 |
| *dmsr-12* | **1.26** | 0.03 | 0.02 | 0.05 | 25.2 | 18.0 |
| *seb-3* | 0.01 | 0.02 | 0.01 | **0.06** | 3.00 | 0.02 |
| *srd-32* | **0.07** | 0.02 | 0.02 | 0.04 | 1.75 | 0.17 |
| *srr-7* | 0.03 | 0.03 | **0.17** | 0.04 | 4.25 | 0.68 |
| *pkd-2* | **1.36** | 0.09 | 0.42 | 1.15 | 1.18 | 0.15 |
| *trf-1* | 0.23 | 0.04 | 0.01 | **78.02** | 339.22 | 8.72 |
| *srw-98* | 0.02 | 0.03 | 0.02 | **0.05** | 1.67 | 0.20 |
| *dmsr-13* | 0.02 | 0.03 | 0.02 | **0.05** | 1.67 | 0.45 |
| *dmsr-10* | 0.02 | 0.04 | 0.03 | **0.07** | 1.75 | 1.40 |
| *dmsr-11* | 0.01 | 0.02 | 0.02 | **0.04** | 2.00 | 0.33 |
| *dmsr-16* | 0.02 | 0.04 | 0.03 | **0.06** | 1.50 | 0.67 |

**Figure 1.**
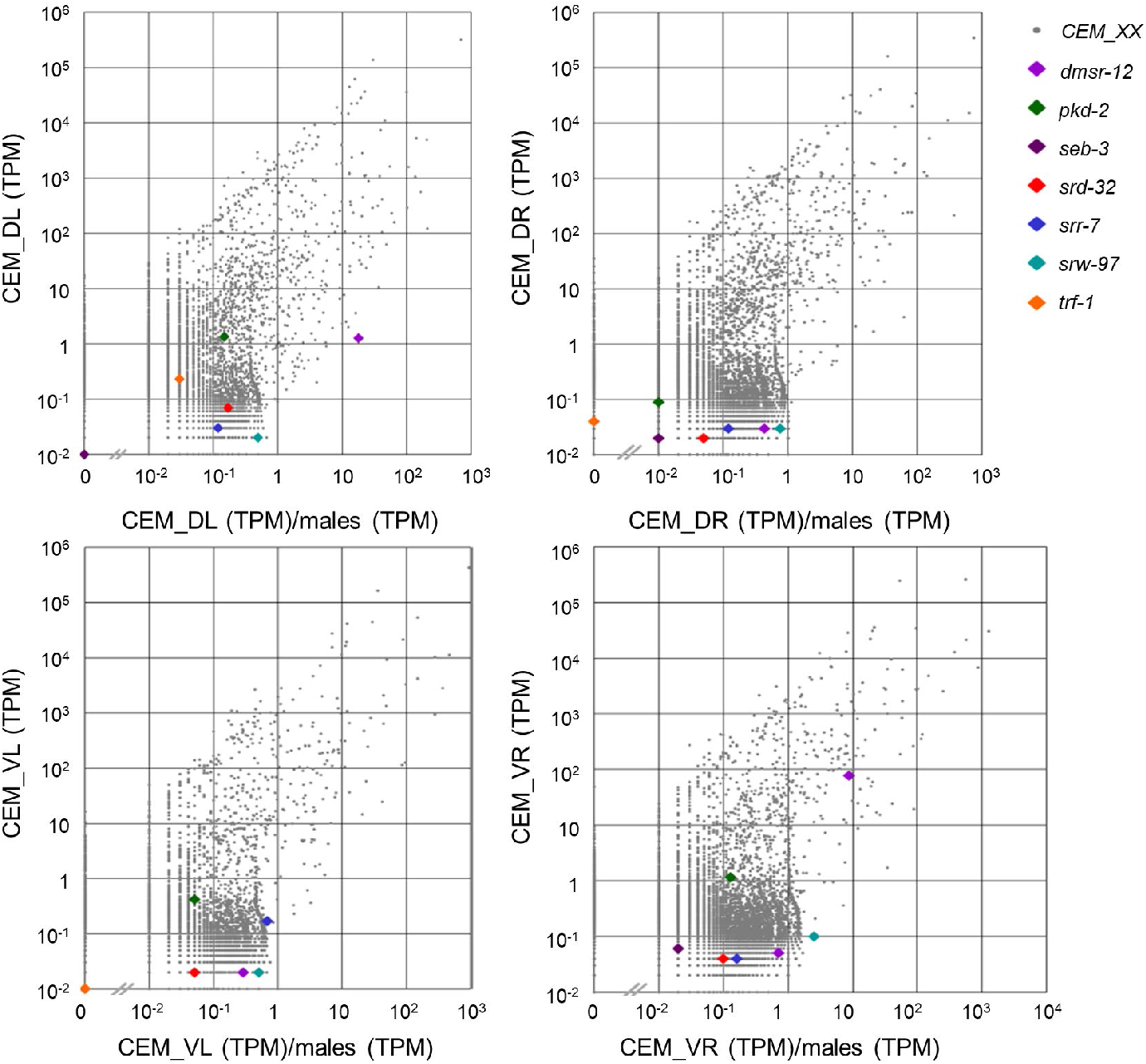
Transcriptomic landscapes of the CEM neurons. **(A-D)** Expression plots of genes expressed in individual CEM DL **(A)**, CEM DR **(B)**, CEM VL **(C)**, and CEM VR **(D)** neurons. For each gene, its X-axis position displays the ratio of its expression in an individual CEM neuron divided by its highest expression in three replicates of adult *C. elegans* males; its Y-axis position displays expression in an individual CEM neuron type. Expression levels are measured in transcripts per million (TPM). Genes of interest are denoted by colored symbols, defined in the legend.

### CEM-specific receptor expression patterns

To confirm our single-cell RNA-seq results, we generated transgenic GFP fusions for the five receptor genes (Boulin *et al*. 2006). Roughly 3 kb of promoter region upstream of the start codon was included in these constructs, along with the majority of the coding sequence (**Table S6**); this would have automatically included any large 5’-ward introns that might contain cis-regulatory elements of these genes (FUXMAN BASS *et al*. 2014). The GFP coding sequence was cloned from the Fire Kit vector pPD95.75 (Boulin *et al*. 2006). *pha-1; lite-1; him-5* animals were injected with reporter constructs and the co-injection marker pBX (*pha-1*(+)). We isolated GFP^+^ strains and imaged GFP^+^ males for expression at 63x magnification (**Figure 2; Fig. S1**). An integrated p*pkd-2*::GFP line was used as a CEM-specific control (**Figure 2A**).

**Figure 2.**
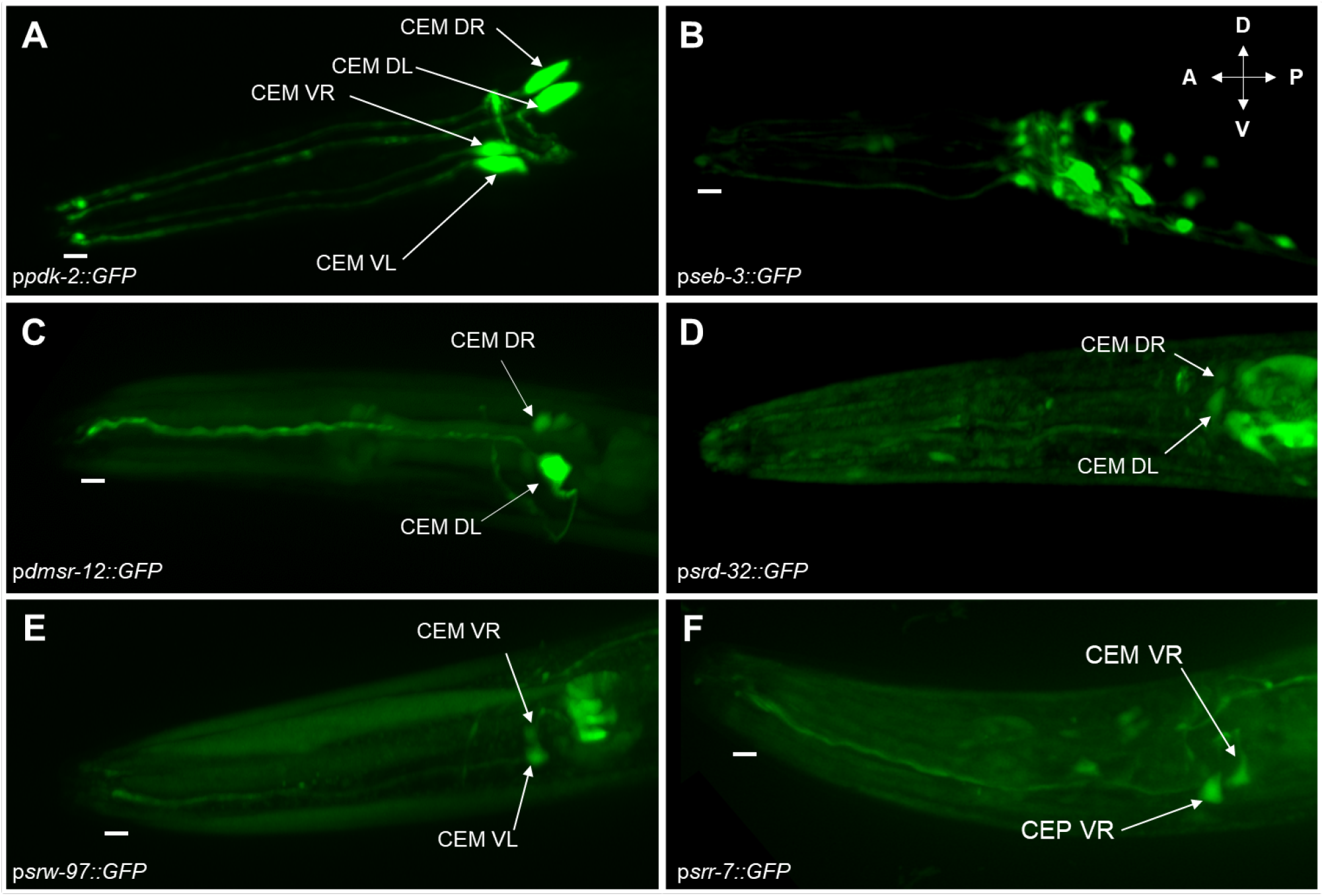
Expression profiles of CEM-enriched GPCR genes of interest. Transcriptional GFP fusions of GPCR genes enriched in the CEM neurons. **(A)** A previously published CEM reporter, p*pkd-2*::GFP. **(B)** p*seb*-3::GFP, matching previously published expression, with no discernable enrichment in the CEM. **(C)** p*dmsr-12*::GFP is strongly expressed in the CEM DL neuron, as well as the CEM DR soma. **(D)** p*srd-32*::GFP is weakly expressed in the somas of CEM DR and DL. **(E)** p*srw-97*::GFP is expressed in both CEM VR and VL, with localization in the cilia as well as the soma. **(F)** p*srr-7*::GFP is expressed in CEM VR, both the soma and cilia, as well the cilial of a neighboring neuron, presumably CEP VR. Dorsal/ventral axes and anterior/posterior directions shown in (B). Scale bars denote distal, cilia region.

The previously characterized GPCR gene, *seb-3*, displayed a non-CEM-specific expression pattern matching that previously described (**Figure 2B**). The other four receptor genes showed transgenic expression patterns similar to their RNA-seq data (**Figure 2C-E**). SRR-7 was observed in CEM VR, as well as another neuron that is likely to be the mechanoreceptor CEP VR (Sawin *et al*. 2000) (**Figure 2F**). Both *seb-3* and *srr-7* were excluded from further analyses, as we aimed to identify CEM-specific regulators of the ascr#8 response.

Except for SRD-32, all GFP-tagged receptors showed subcellular localization patterns that included the sensory cilia (**Figure 2, white bars**). Our failure to see this for SRD-32 may be an artifact of our p*srd-32*::GFP construct design, because only 52% of the *srd-32* coding sequence was included in our transgene (**Table S6**). We also observed non-GPCR genes to be specifically expressed in single CEM neurons, such as *trf-1*, which encodes a TNF receptor homolog (Tenor and Aballay 2008) expressed in CEM VR (**Table 1**); *trf-1* has an EVN-specific promoter (Wang *et al*. 2015), providing an alternative to the *pkd-2* and *klp-6* promoters typically used to drive transgenes in EVNs (**Fig. S1**) (Peden and Barr 2005; Bae *et al*. 2006).

### RNAi-mediated knockdown of CEM Receptors

To test whether these CEM-enriched genes encode receptors that are required in ascr#8 sensation, we fed dsRNA to a strain that is hypersensitive to neuronal RNA interference, *nre-1; lin-15B* (Schmitz *et al*. 2007; Poole *et al*. 2011).

To confirm that our system for male neuronal RNAi could affect behavioral phenotypes, we first fed animals *osm-3* and *osm-9* dsRNA clones from the Ahringer Library (Fraser *et al*. 2000; Kamath *et al*. 2003). Using a Spot Retention Assay (Narayan *et al*. 2016), we assayed animals for their behavioral dwell time in ascr#8 (**Figure S2A**). Animals fed *osm-3* dsRNA showed significantly defective responses to ascr#8, as observed for loss-of-function alleles (**Figure S2B**). In contrast, animals fed *osm-9* showed only a slight decrease in their times spent within ascr#8 (**Figure S2A**). Because we could successfully abolish ascr#8 attraction through male neuronal RNAi of *osm-3*, we were confident that RNAi would be effective at functionally verifying CEM-enriched genes encoding components of the ascr#8 response.

We fed animals dsRNA clones targeting three CEM-enriched receptor genes: *srd-32, dmsr-12*, and *srw-97*. A *srd-32* clone was unavailable in the Ahringer library (Fraser *et al*. 2000; Kamath *et al*. 2003), but a present in the Vidal library (Rual *et al*. 2004). Each library uses the same backbone vector, allowing the same control to be used for both.

RNAi of *srd-32* caused no defect in ascr#8 responses (**Figure S2C**). Both *dmsr-12* or *srw-* 97 dsRNA caused partial defects of ascr#8 dwell time (**Figure S2C**). Neither knockdown statistically lowered the dwell time in ascr#8 (**Figure S2D**), though they did abolish the statistically significant increase in time spent in ascr#8 over vehicle controls (**Figure S2C**).

### CRISPR-generated null mutants of candidate receptors

For phenotypic analysis of null mutants, we used our Single Worm Attraction Assay (SWAA), as previously described (Reilly *et al*. 2021). Wild-type (*him-5*) animals were strongly attracted to ascr#8 in the SWAA, replicating previous observations (**Figure 3A, B; Fig. S3**) (Reilly *et al*. 2021).

**Figure 3.**
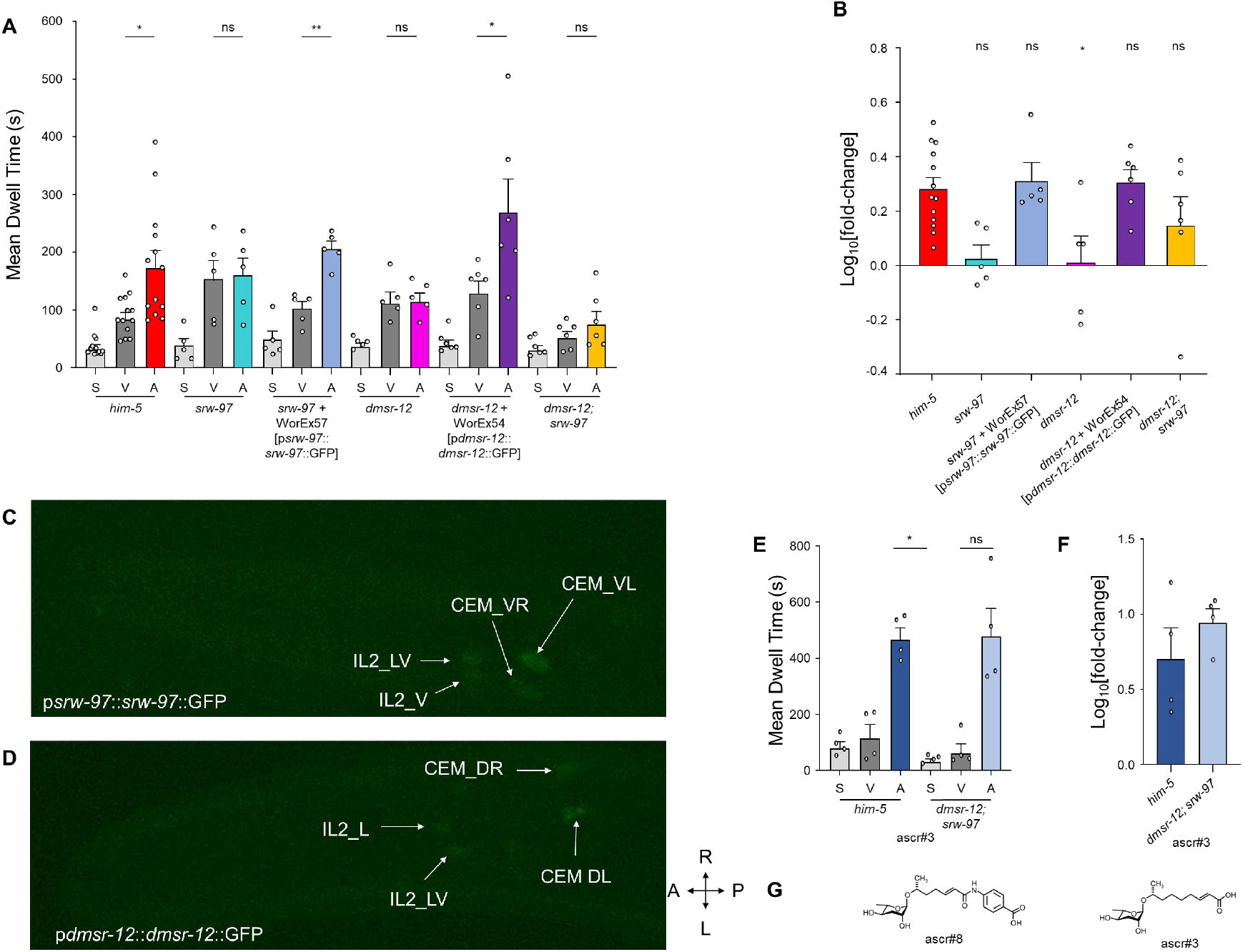
The GPCRs, SRW-97 and DMSR-12, act in non-overlapping CEM neurons to aid in driving the ascr#8 response. **(A)** Raw dwell times in the centers of spatial control (S), vehicle-only control (V), and ascaroside (A) wells, and **(B)** Log(fold-change) of A versus V for *srw-97* knockout and transgenic rescue (WorEx57), *dmsr-12* and transgenic rescue (WorEx54), as well as *srw-97*;*dmsr-12* double mutant animals. **(C)** Expression of WorEx61 (p*srw-97*::*srw-97*::GFP) in CEM VR and CEM VL. **(D)** Expression of WorEx60 (p*dmsr-12*::*dmsr-12*::GFP) in CEM DL and CEM DR. IL2_LV and IL2_L somas denoted. Cellular identities confirmed through co-expression of p*klp-6*::dtTomato (**Fig. S5**). Anterior/Posterior and Right/Left axes denoted. **(E)** Raw dwell times and **(F)** Log(fold-change) of *him-5* and *dmsr-12;srw-97* animals in response to the structurally related attracted pheromone, ascr#3. **(G)** Structural comparison of ascr#8 (left) and ascr#3 (right). Error bars denote SEM. *n* ≥ 5. **(A, E)** RM-ANOVA followed by Dunnett’s post-hoc test or Friedman test followed by Dunn’s (within strain), **(B)** ANOVA followed by Bonferroni’s post-hoc test (between strain log(fold-change) values). **(F)** Student’s t-test between strain log(fold-change) values * *p* < 0.05, ** *p* < 0.01.

*srw-97(knu456)* males displayed partially defective ascr#8 attraction: their ascr#8 dwell time was no different than that of vehicle, nor was it different that *him-5* animals (**Figure 3A, B; Fig. S3)**. We were able to restore normal ascr#8 attraction in *srw-97(knu456)* males through transgenesis with a translational fusion construct (p*srw-97*::*srw-97*::GFP) (**Fig. S4**). Expression of the rescue transgene matched that of our initial fusion, with GFP visible in both ventral CEM neurons (**Figure 3C; Fig. S5**).

Unlike *srw-97(knu456), dmsr-12(tm8706)* males exhibited completely defective responses to ascr#8 (**Figure 3A, B; Fig. S3**). Similarly, the expression profile of the p*dmsr-12*::*dmsr-12*::GFP rescue construct (**Fig. S4**) matched its earlier reporter expression (**Figure 3D; Fig. S5**), and completely restored attraction to ascr#8 (**Figure 3A, B; Fig. S3**). Quantification of GFP expression confirmed SRW-97 to be enriched within the CEM_VL neuron, and to a lesser extent CEM_VR and CEM_DL neurons, while DMSR-12 was observed within CEM_DL and CEM_DR neurons (**Table S7**).

Given that *dmsr-12* is expressed in dorsal CEM neurons, and *srw-97* is expressed in ventral CEM neurons (**Figure 3C, D**), we speculated that single mutants of either receptor partially retain the ability to sense and respond to ascr#8, with the remaining CEM-specific receptor gene conferring some residual response to ascr#8. We thus generated a *dmsr-12; srw-97* double mutant strain and assayed animals for their ability to respond to ascr#8. These double mutants were not attracted to ascr#8 compared to the vehicle control (**Figure 3A; Fig. S3**), though they were not significantly defective in their attraction compared to wild-type animals either (**Figure 3B**). However, this is likely a result of lower vehicle dwell times in the double mutants. This defect was specific to ascr#8; double mutants showed no defect in their responses to ascr#3 (**Figure 3E, F; Fig. S6**).

Together, these data suggest that at least two GPCR receptors are expressed in discrete pairs of CEM neurons and act in parallel during sensation of ascr#8.

### Phylogenetic analyses of ascr#8 receptors reveal likely gene duplication events

The ability to attract mates through pheromones is often essential for a species’ survival. We have recently observed that different *Caenorhabditis* species show variable levels of attraction to ascr#8 (Reilly *et al*. 2019). To better understand variability in levels of attraction, we analyzed the evolution of both *srw-97* and *dmsr-12*.

We identified orthologs of *srw-97* from *C. elegans* (*Cel-srw-97*) in other *Caenorhabditis* proteomes via OrthoFinder (**Figure 4A**). A closely related *C. elegans* paralog, *Cel-srw-98*, underwent a species-specific expansion within *C. inopinata*. The coding DNA sequences (CDS DNAs) of both *Cel-srw-97* and *Cel-srw-98* are 66.2% identical to one another, while their amino acid sequences are only 57.1% identical (Madeira *et al*. 2019). *Cel-srw-98* shows enriched expression in CEM VR that is similar to (though weaker than) that of *Cel-srw-97* (**Table 1; Table S5**).

**Figure 4.**
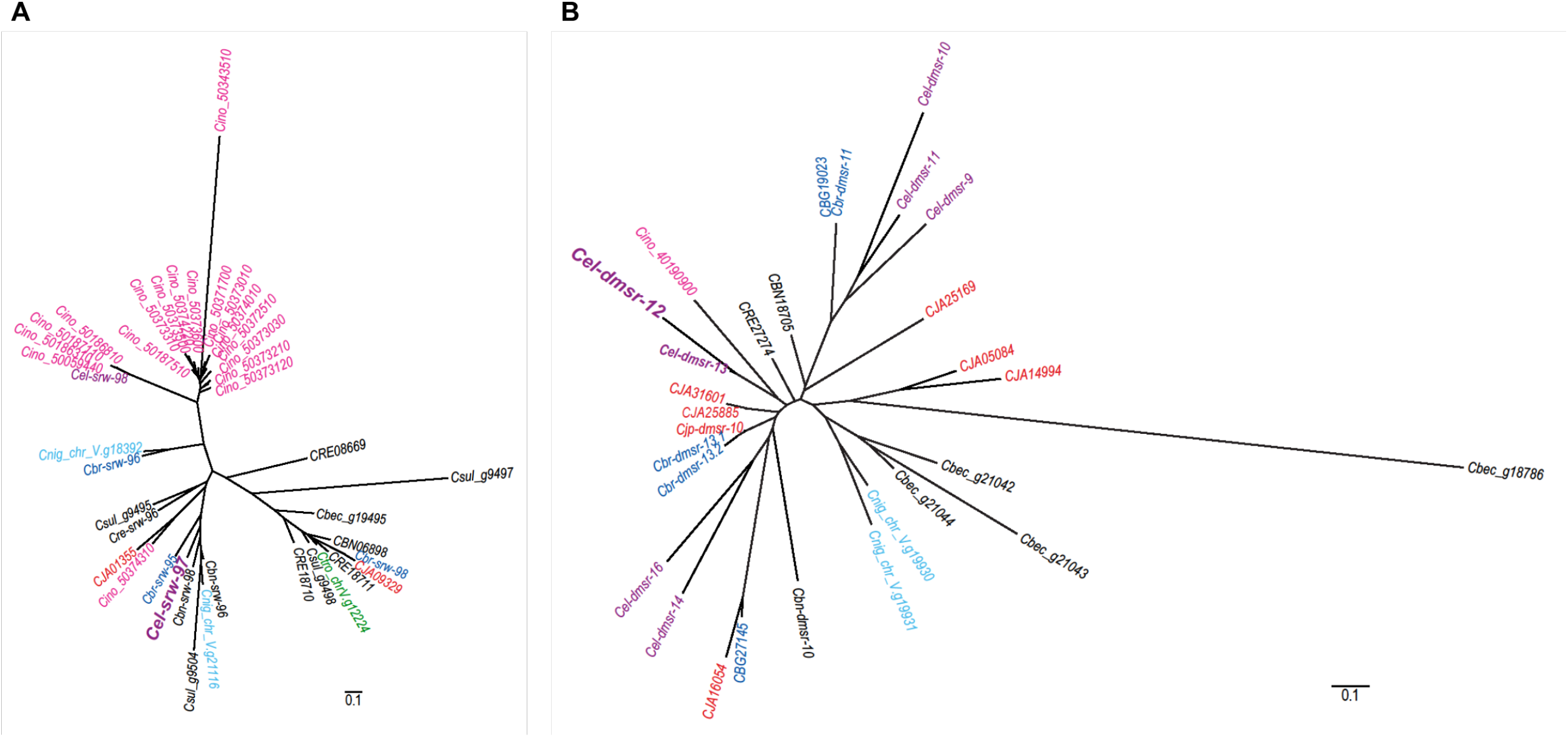
Phylogenetic analysis of ascr#8-receptor candidate paralogs across the *Caenorhabditis* genus. **(A)** The phylogeny of *Cel_srw-97* reveals a *C. inopinata*-specific amplification of *Cel_srw-98*. **(B)** The phylogeny of *Cel_dmsr-12* shows conserved numbers of orthologs across the genus. Genes for *C. elegans* are denoted in purple, *C. inopinata* in pink, *C. nigoni* in light blue, *C. briggsae* in dark blue, *C. japonica* in red, and *C. tropicalis* in dark green. Distance reference bars (0.1) depict substitutions per site.

*C. tropicalis*, like *C. elegans* and *C. briggsae*, is a hermaphroditic *Caenorhabditis* species (Noble *et al*. 2021) which encodes a reduced number of ascr#8 receptor gene paralogs, with no *Cel-dmsr-12* paralogs, and only one *Cel-srw-97* paralog (**Figure 4**). We previously observed that *C. tropicalis* fails to be attracted to ascr#8, and it is possible that this loss of receptor genes is one reason for that failure (Reilly *et al*. 2019).

In contrast, the other candidate ascr#8 receptor, *dmsr-12*, does not appear in our phylogenetic analysis to have undergone any species-specific expansion (**Figure 4B**). Rather, there are generally fewer paralogs per non-*C. elegans* species.

## Discussion

Pheromones are important for mating in many animals. In the nematode *C. elegans*, males are attracted to hermaphrodites as possible mates by the small molecule pheromone ascaroside #8 (Pungaliya *et al*. 2009; Narayan *et al*. 2016). While the ascaroside class of small molecule pheromones utilized by nematodes is rapidly being elucidated (with over 230 known ascaroside structures so far; *https://smid-db.org*) (Artyukhin *et al*. 2018), the neuronal receptor proteins that mediate pheromone signals remain largely unknown. For only a select few ascarosides have sensory components been identified at the cellular (Greene *et al*. 2016a; Greene *et al*. 2016b; Chute *et al*. 2019), receptor (Kim *et al*. 2009; Mcgrath *et al*. 2011; Park *et al*. 2012), or signal transduction levels (Zwaal *et al*. 1997).

Here, we identify two novel G protein-coupled receptors as active, required components in sensory and behavioral responses to the mating pheromone ascr#8. Transcripts for the two GPCRs, *dmsr-12* and *srw-97*, are enriched in single CEM neurons (**Table 1**), and they express in non-overlapping subsets of the male-specific chemosensory neurons (**Figure 1, 2**). There are, however, other receptors present in these same neurons that contribute to the navigation of a vast and ever-changing array of environmental cues, such as the widely expressed ethanol sensor *seb-3* (**Figure 2B**) (Jee *et al*. 2013).

One mechanism for sensory flexibility in CEM neurons may be heterodimerization of receptor proteins. Previous work identified two receptors for ascr#2 in the ASK neuron: DAF-37 and DAF-38 (Park *et al*. 2012). While both are required for proper ascr#2-induced dauer formation, only DAF-38 is involved in the sensation of other ascarosides (Park *et al*. 2012). Such heterodimers may also exist for pheromone receptors in CEM neurons.

Another mechanism for sensory flexibility in CEM neurons is to have diverse receptor proteins with distinct specificities co-expressed within a single neuron. Multiple ascarosides are sensed by the male-specific CEM neurons, including ascr#8 and ascr#3 (Narayan *et al*. 2016). The two receptors identified here may function as ascr#8-specific receptors, as *dmsr-12;srw-97* double mutant animals do not exhibit defective responses to ascr#3 (**Figure 3E, F**). The related receptors *dmsr-10, dmsr-13, dmsr-16*, and *srw-98* are enriched in ventral CEM neurons, and may heterodimerize with *srw-97*, to result in fine-tuned sensation of ascr#8. Similarly, there may be other receptors may be enriched in the dorsal CEM neurons to heterodimerize with *dmsr-12*. The role of DAF-37 as an ascr#2-specific receptor further supports our hypothesis, as *dmsr-12* is related to DAF-37 (Robertson and Thomas 2006). The ventral CEM receptor SRW-97 falls within the same large family of GPCRs as well (Krishnan *et al*. 2014), while the promiscuous DAF-38 does not (Robertson and Thomas 2006).

The initial goal of our single-cell RNA-seq analysis of male-specific chemosensory CEM neurons was to identify GPCRs encoding pheromone receptors. However, further analysis of other genes with CEM-enriched expression should also uncover specific and novel promoter profiles, which will enable optogenetic manipulation and calcium imaging of defined individual CEM neurons (Narayan *et al*. 2016). The development of advanced transgenic reagents will also permit chemical biology of CEM neurons. We have recently developed an active ascr#8 bioaffinity probe to employ in the targeted elucidation of ascr#8 receptors (Zhang *et al*. 2019). Use of this probe will lead to further elucidation of the identity of acsr#8 receptors, by confirming either SRW-97 or DMSR-12 as receptors or identifying their heterodimeric partners. The combination of these technologies should clarify how heterogeneous CEM neurons achieve homogeneous sensory responses.

## Methods

### Single cell isolation and RT-PCR

Individual ciliated male-specific cephalic sensory (CEM) neurons were isolated from adult males of the *C. elegans* strain CU607 (smIs23 [p*pkd-2*::GFP+ pBX]; *him-5*(e1490)), as previously described (Narayan *et al*. 2016). This strain expresses an integrated GFP transgene that labels male-specific extracellular vesicle-releasing neurons (EVNs, p*pkd-2*::GFP [Figure 2A]). Cells were separated by anatomical identity (i.e., CEM dorsal left [CEM_DL], CEM dorsal right [CEM_DR], CEM ventral left [CEM_VL], and CEM ventral right [CEM_VR]). Microdissection and single-cell RT-PCR of individual CEM_DL, CEM_DR, CEM_VL, and CEM_DR neurons was performed essentially as described (Schwarz *et al*. 2012). For all four neuronal types, single-end 50-nucleotide (50-nt) to 100-nt RNA-seq was performed on an Illumina HiSeq 2000. To identify so-called housekeeping genes and genes primarily active outside the nervous system, we compared our results from CEM_DL, CEM_DR, CEM_VL, and CEM_DR to equivalent reanalyses of published single-end 38-nt RNA-seq data from mixed-stage whole *C. elegans* hermaphrodite larvae (Schwarz *et al*. 2012), and of published single-end 36-nt RNA-seq data from adult whole *C. elegans* males and females (Thomas *et al*. 2012).

### Transcriptional analysis

Reads from RT-PCR-amplified CEM neurons and larvae were quality-filtered as follows: neuronal reads that failed Chastity filtering were discarded (Chastity filtering had not been available for the larval reads); raw 38-nt larval reads were trimmed 1 nt to 37 nt; all reads were trimmed to remove any indeterminate (“N”) residues or residues with a quality score of less than 3; and larval reads that had been trimmed below 37 nt were deleted, as were neuronal reads that had been trimmed below 50 nt. For uniformity of analysis, all RNA-seq reads of 51-100 nt in length were trimmed to 50 nt before further RNA-seq analysis. This left a total ranging from 55,902,297 (CEM_VL) to 73,412,777 (CEM_VR) filtered reads for analysis of each neuronal type, versus 23,369,056 filtered reads for whole larvae (**Table S2**). Previously published RNA-seq reads (Thomas *et al*. 2012) from three biological replicates of adult males (accession numbers SRR580386, SRR580387, and SRR580388) and adult females (accession numbers SRR580383, SRR580384, and SRR580385) were downloaded from the European Nucleotide Archive (*ftp://ftp.sra.ebi.ac.uk*) and used without further trimming or filtering.

We used RSEM version 1.2.17 (Li and Dewey 2011) with bowtie2 version 2.2.3 (Langmead and Salzberg 2012) and SAMTools version 1.0 (Li *et al*. 2009) to map filtered reads to a *C. elegans* gene index and generate read counts and gene expression levels in transcripts per million (TPM). To create the *C. elegans* gene index, we ran RSEM’s *rsem-prepare-reference* with the arguments ‘‘*--bowtie2 --transcript-to-gene-map*’ upon a collection of coding DNA sequences (CDSes) from both protein-coding and non-protein-coding *C. elegans* genes in WormBase release WS245 (Howe *et al*. 2016). The CDS sequences were obtained from *ftp://ftp.sanger.ac.uk/pub2/wormbase/releases/WS245/species/c_elegans/PRJNA13758/c_elegans.PRJNA13758.WS245.mRNA_transcripts.fa.gz* and *ftp://ftp.sanger.ac.uk/pub2/wormbase/releases/WS245/species/c_elegans/PRJNA13758/c_elegans.PRJNA13758.WS245.ncRNA_transcripts.fa.gz*. For each RNA-seq data set of interest, we computed mapped reads and expression levels per gene by running RSEM’’s *rsem-calculate-expression* with the arguments ‘‘*--bowtie2 -p 8 --no-bam-output --calc-pme --calc-ci --ci-credibility-level 0.99 --fragment-length-mean 200 --fragment-length-sd 20 --estimate-rspd --ci-memory 30000*’. These arguments, in particular ‘*--estimate-rspd*’, were aimed at dealing with single-end data from 3’-biased RT-PCR reactions; the argument ‘*--phred64-quals*’ was used for the larval reads, while ‘*--phred33-quals*’ was used for all other reads (neuronal, adult males, and adult females). We computed posterior mean estimates (PMEs) both for read counts and for gene expression levels, and rounded PMEs of read counts down to the nearest lesser integer. We also computed 99% credibility intervals (Cis) for expression data, so that we could use the minimum value in the 99% CI for TPM as a robust minimum estimate of a gene’s expression (minTPM).

We observed the following overall alignment rates of reads to the WS245 *C. elegans* gene index: 47.16% for the CEM_DL read set, 27.37% for the CEM_DR read set, 40.20% for the CEM_VL read set, 17.99% for the CEM_VR read set, and 76.41% for the larval read set (**Table S2**). A similar discrepancy between lower alignment rates for hand-dissected linker cell RNA-seq reads versus higher alignment rates for whole larval RNA-seq reads was previously observed, and found to be due to a much higher rate of human contaminant RNA sequences in the hand-dissected linker cells (Schwarz *et al*. 2012). For previously published adult male and female reads, we observed mapping rates of 88.20% to 96.11%; this may reflect greater difficulty of getting mappable reads from our RT-PCR-amplified cDNAs than from unamplified cDNA syntheses.

We defined detectable expression for a gene in a given RNA-seq data set by that gene having an expression level of 0.1 TPM; we defined robust expression by that gene having a minimum estimated expression level (termed minTPM) of at least 0.1 TPM in a credibility interval of 99% (i.e., ≥ 0.1 minTPM). The numbers of genes being scored as expressed in a given neuronal type above background levels, for various data sets, are given in **Table S3**. Other results from RSEM analysis are given in **Table S5**.

We annotated *C. elegans* genes and the encoded gene products in several ways (**Table S5**). For the products of protein-coding genes, we predicted classical signal and transmembrane sequences with Phobius 1.01 (KÄLL *et al*. 2004), regions of low sequence complexity with pseg (SEG for proteins, from *ftp://ftp.ncbi.nlm.nih.gov/pub/seg/pseg*) (WOOTTON 1994), and coiled-coil domains with ncoils (from *http://www.russell.embl-heidelberg.de/coils/coils.tar.gz*) (LUPAS 1996). Protein domains from Pfam 31.0 (Finn *et al*. 2016) were detected with HMMER 3.1b2/ *hmmsearch* (EDDY 2009), using the argument ‘--*cut_ga*’ to invoke model-specific reporting thresholds. Memberships of genes in orthology groups from eggNOG 3.0 (Powell *et al*. 2012) were extracted from WormBase WS245 with the TableMaker function of ACEDB 4.3.39. Genes with likely housekeeping status (based on ubiquitous expression in both larvae and linker cells) were as identified in our previous work (Schwarz *et al*. 2012). Genes were predicted to encode GPCRs on the basis of their encoding a product containing one or more of the following Pfam-A protein domains: 7tm_1 [PF00001.16], 7tm_2 [PF00002.19], 7tm_3 [PF00003.17], 7tm_7 [PF08395.7], 7TM_GPCR_Srab [PF10292.4], 7TM_GPCR_Sra [PF02117.11], 7TM_GPCR_Srbc [PF10316.4], 7TM_GPCR_Srb [PF02175.11], 7TM_GPCR_Srd [PF10317.4], 7TM_GPCR_Srh [PF10318.4], 7TM_GPCR_Sri [PF10327.4], 7TM_GPCR_Srj [PF10319.4], 7TM_GPCR_Srsx [PF10320.4], 7TM_GPCR_Srt [PF10321.4], 7TM_GPCR_Sru [PF10322.4], 7TM_GPCR_Srv [PF10323.4], 7TM_GPCR_Srw [PF10324.4], 7TM_GPCR_Srx [PF10328.4], 7TM_GPCR_Srz [PF10325.4], 7TM_GPCR_Str [PF10326.4], ABA_GPCR [PF12430.3], Sre [PF03125.13], and Srg [PF02118.16]. By this criterion, we identified 1,615 genes encoding GPCRs in the WS245 version of the *C. elegans* genome; this resembles a previous estimate of ∼1,470 *C. elegans* genes encoding chemoreceptors and other GPCRs, identified through extensive computational and manual analysis (HOBERT 2013). The memberships of genes in orthology groups from eggNOG 3.0 (Powell *et al*. 2012) were extracted directly from WormBase WS245 with the TableMaker function of ACEDB 4.3.39. Genes with likely housekeeping status (based on ubiquitous expression in both larvae and linker cells) were as identified in our previous work (Schwarz *et al*. 2012). Gene Ontology (GO) annotations for *C. elegans* genes were extracted from WormBase-computed annotations in *ftp://ftp.wormbase.org/pub/wormbase/releases/WS245/ONTOLOGY/gene_association.WS245.wb.c_elegans*; human-readable text descriptions for GO term IDs were extracted from *term.txt* in the Gene Ontology archive *http://archive.geneontology.org/full/2014-07-01/go_201407-termdb-tables.tar.gz*.

### GFP reporter construction

Reporter fusion constructs were generated using previously described techniques (Boulin *et al*. 2006). Approximately 2-3 kb of upstream promoter region of each gene was included in construct generation, as well as a portion of the coding sequence (**Table S6**). This was then fused to GFP (from the Fire Vector Kit plasmid, pPD95.75), via PCR fusion (Boulin *et al*. 2006). Primers were designed using Primer 3 and ordered from IDT (Integrated DNA Technologies). Primer sequences available in **Table S9**. Successful fusion was confirmed via gel electrophoresis prior to injection.

Reporter fusion constructs were injected into the gonads of *pha-1(e2123ts); lite-1(ce314); him-5(e1490)* animals, along with a co-injection marker of pBX(*pha-1*(+)). In this manner, positive array animals will propagate normally at 20 °C. Strains were confirmed via GFP expression, with multiple array lines being generated per injection (**Table S8**). Injections were performed by InVivo Biosystems, with strain isolation being performed in house.

### Imaging

Animals were imaged for GFP expression using previously described techniques. GFP+ young adult male animals were mounted on a 1% agarose pad and immobilized with sodium azide. Animals were then imaged on a spinning disk confocal microscope at 63x magnification. Z-stack imaging was performed, generating 3D reconstructions of the heads of the imaged animals. Central/optimal z-plane images were used to generate the images used to verify expression (**Figure 2; Fig. S1**).

### RNAi Feeding

VH624 (*rhIs13* [*unc-119*::GFP + *dpy-20*(+)]; *nre-1(hd20)*; *lin-15B(hd126)*) animals (Schmitz *et al*. 2007; Poole *et al*. 2011) were crossed with *him-5(e1490)* animals to integrate male production into a strain hypersensitive to neuronal RNAi-knockdown, generating JSR44 (*nre-1(hd20)*; *lin-15B(hd126); him-5(e1490)*). During the cross, insertion of the *him-5(e1490)* allele displaced the integrated array *rhIs13*, suggesting location of the array on Chromosome V. Presence of *lin-15B(hd126)* in JSR44 was confirmed via sequencing. The non-annotated *nre-1(hd20)* is linked with *lin-15B*, being retained alongside *lin-15B* (Schmitz *et al*. 2007; Poole *et al*. 2011).

RNAi clones were grown overnight in cultures of LB containing 50 μg/mL ampicillin. Cultures were then diluted to an OD600 of 1.0 before plating on NGM agar plates containing 50 μg/mL ampicillin and 1 mM IPTG (Isopropyl-β-D-thiogalactoside) to select for RNAi clones and induce expression. Lawns were allowed to grow at room temperature for 8-16 hours, before JSR44 young adult hermaphrodites were placed on the plates and left to propagate at 16°C. Young adult males of the F1 progeny were then selected for behavioral testing suing the *Spot Retention Assay*. Empty vector controls (VC-1 clone) were run alongside every targeted knockdown experiment.

### Spot Retention Assay

Following previously described methods, young adult males were isolated from hermaphrodites 5-16 hours prior to testing (Pungaliya *et al*. 2009; Narayan *et al*. 2016). In short, at the time of the assay, 0.6 μL of either vehicle control (-) or ascaroside #8 (+) was added to the NGM plates covered in a thin lawn of OP50 *E. coli*. Ten males were then divided between two pre-marked spots on the agar, equidistant from the cues. The plate was then recorded for 20 minutes. The time spent of each visit in either vehicle or ascaroside #8 (if greater than 10 seconds) was scored and averaged. Plates in which the average was greater than two standard deviations removed from the population average were removed from the final analysis as outliers. To compare between strains or conditions, the vehicle was subtracted from the ascaroside dwell time for each plate. The average of these differences was then compared statistically.

### Single Worm Behavioral Assay

Following previously described methods (Pungaliya *et al*. 2009; Narayan *et al*. 2016; Reilly *et al*. 2021), animals were isolated and prepared in an identical manner to the Spot Retention Assay. The two outside rings of wells in a 48-well tissue culture plate were filled with 200 μL of NGM agar, which was then seeded with 65 μL of OP50 *E. coli*. The plates and lawns were then dried at 37 °C for 4 hours. Alternating wells were then prepared as spatial controls (“S”, nothing done), vehicle controls (“V”, 0.85 μL of dH2O was placed in the center of the well), or ascaroside well (“A”, 0.85 μL of ascaroside was placed in the center of the well). This was performed over four quadrants. Animals were scored for their visits and duration to the center of the well and/or the cue.

The average duration of each worm’s visits was calculated, and these values were again averaged together to generate a Mean Dwell Time in seconds for each plate. When comparing across strains or conditions, the Spatial controls were then compared for statistical difference. If none was observed, the Log(fold-change) A/V was then calculated by taking the log of the ascaroside mean dwell time divided by the vehicle mean dwell time for each plate. The number of times each worm visited the center was averaged to generate the Visit Counts.

The Percent Attraction values were calculated by first determining the “attractive” cut off as two standard deviations above the vehicle average. Any visit longer than this was deemed “attractive” and scored as a “1”; non-attractive visits were scored as “0”. The percent attraction was then calculated for each worm was the percent of visits scored a “1”. The average was then calculated across the plate to determine Percent Attraction.

### CRISPR Design and Strain Generation

A novel null mutation was generated for *srw-97* by InVivo Biosystems. The *srw-97(knu456)* allele was generated in a *him-5(e1490)* strain using two sgRNAs (TTTAGTAGAGCAGAAATTAA and TACAGCTTTAACTTTCAAC) to generate a 1,620-nt deletion which removed the start codon and left only the terminal exon intact. The *knu456* knockout allele was generated by donor homology using the pNU1361odn oligo: (TTTTCTTGTATTTCCAAAAATTGTAAAAACCTTTATGAAAGTTAAAGCTGTAAGGATTTTCAGACATTTA). Following generation of a homozygous deletion by InVivo Biosystems, the line was then backcrossed twice into *him-5(e1490)*.

The *dmsr-12(tm8706)* allele, provided by the National BioResource Group (NRBP) contains a 118-nucleotide deletion that was generated by Dr. Mitani of the NRBP. The allele was crossed into a *him-5(e1490)* background prior to testing. The deletion spans intron 2 and exon 3 of the coding sequence. Whether this results in a correctly spliced gene remains unknown, although the expected remaining coding sequence remains in frame.

### Phylogenetic Analyses

For phylogenetic analysis of selected CEM genes, we downloaded proteomes for *C. elegans* and related *Caenorhabditis* nematodes from WormBase (release WS275), the Blaxter *Caenorhabditis* database (release 1), or our unpublished work, as listed in **Table S4**. From each proteome, we extracted the longest predicted isoform for each gene with *get_largest_isoforms.pl* (*https://github.com/SchwarzEM/ems_perl/blob/master/fasta/get_largest_isoforms.pl*). We observed that the predicted isoform for *dmsr-12* in the WormBase WS275 release of *C. elegans*’ proteome was shorter than past versions of *dmsr-12*, and that the WS275 isoform omitted exons that our transgenic expression data (based on older gene models for *dmsr-12*) indicated were likely to be real. We therefore manually replaced the WS275 version of *dmsr-12* with an older version (extracted from the *C. elegans* proteome in the WS250 release of WormBase). We then computed orthology groups for *C. elegans* and its related species with OrthoFinder version 2.3.11 (Emms and Kelly 2015; Emms and Kelly 2019), using the arguments ‘*-a 1 -S diamond -og*’. We identified which orthology groups contained the *C. elegans* genes *srr-7, srw-97*, and *dmsr-12*, and extracted their sequences from a concatenation of all 11 proteomes via *extract_fasta_subset.pl* (*https://github.com/SchwarzEM/ems_perl/blob/master/fasta/extract_fasta_subset.pl*). For each orthogroup’s member sequences, we aligned the sequences with MAFFT version 7.455 (Katoh and Standley 2013) and filtered the alignments twice with trimAl version 1.4.rev15 (CAPELLA-GUTIÉRREZ *et al*. 2009), using first the argument ‘*-automated1*’ and then the arguments ‘*-resoverlap 0.50 -seqoverlap 50*’. From the filtered alignments, we computed maximum-likelihood protein phylogenies with IQ-TREE version 2.0-rc1 (Nguyen *et al*. 2015; Kalyaanamoorthy *et al*. 2017), using the arguments ‘*-m MFP -b 100 --tbe*’. In particular, we used transfer bootstrap expectation (‘*--tbe*’) which provides more reliable confidence values than classic bootstrapping (Lemoine *et al*. 2018). We visualized the resulting phylogenies with FigTree version 1.4.4 (*http://tree.bio.ed.ac.uk/software/figtree*).

## Statistical Analyses

Prior to any statistical analyses, outliers were identified and removed. Outliers were defined as any data points greater than two standard deviations removed from the average. All data were then tested for normality using a Shapiro-Wilk Normality Test. This test was chosen over the more conventional D’Agostino-Pearson Normality Test as many data sets were below 10 in number (due to the statistical power offered by the Single Worm Behavioral Assay (Reilly *et al*. 2021).

The Spot Retention Assay data was analyzed using paired *t*-tests or Wilcoxon Matched-Pairs Signed Rank tests to compare vehicle control and ascaroside dwell times within strains following tests for normality (**Figure S2**). When comparing the values of multiple conditions or strains, the data was first normalized to account for vehicle dwell time variation between plates using a base-2 exponentiation, as described previously, to transform all data points into non-zero values. This allows for the calculation of the fold-change as the log(base2) of the ascaroside dwell time divided by the vehicle dwell time. These normalized values were then compared using a Mann-Whitney test or a One-Way ANOVA followed by a Dunnett’s multiple comparisons test (**Figure S2**).

The Single Worm Behavioral Assay was first analyzed within each strain by performing either a Repeated-Measures ANOVA followed by a Dunnett’s multiple comparisons test or Friedman test followed by Dunn’s correction, comparing both Spatial control and Ascaroside dwell times to that of the Vehicle Control. The log(fold-change) of raw ascaroside and vehicle dwell time values were calculated and compared using either a One-Way ANOVA followed by a Bonferroni’s multiple corrections test or Friedman Test followed by Dunn’s correction (**Figure 3**). The ascr#3 log(fold-change) values were compared using a Student’s *t*-test (**Figure 3F**). Visit Counts were compared in the same manner as the Mean Dwell Time data, while the Percent Attraction data was analyzed using paired *t*-tests or Wilcoxon Matched-Pairs Signed Rank tests to compare the attractive values of the vehicle and ascaroside (**Figures S3, S4, S5**).

## Data availability

RNA-seq reads have been deposited in the NCBI Sequence Read Archive (SRA) under the BioProject accession number PRJNA781271.

## Acknowledgements

We thank the *Caenorhabditis* Genetics Center, which is funded by the NIH Office of Research Infrastructure Programs (P40 OD01044), as well as the National BioResource Project, Ding Xue (University of Colorado, Boulder), L. Rene Garcia (Texas A&M University), and Douglas Portman (University of Rochester Medical Center) for providing strains. We also thank Igor Antoshechkin, Caltech Genomics facility for sequencing and InVivo Biosystems for generating transgenic and CRISPR knockout animals. The synthetic ascr#8 utilized in this study was provided by Frank Schroeder (Cornell University). The research reported in this publication was supported by NIH R01 DC016058 (J.S.), R01 GM084389 (P.W.S.), the Howard Hughes Medical Institute (P.W.S.), Moore Foundation Grant No. 4551 (E.M.S.), and Cornell startup funding (E.M.S.). We thank Titus Brown and the Michigan State University High-Performance Computing Center (supported by U.S. Department of Agriculture grant 2010-65205-20361 and NIFA–National Science Foundation (NSF) grant IOS-0923812) for computational support; additional computing was enabled by start-up and research allocations from NSF XSEDE (TG-MCB180039 and TG-MCB190010).

