## Supplemental Text for "Transcriptomic profiling of sex-specific olfactory neurons reveals subset-specific receptor expression in *Caenorhabditis elegans*"

[1]Department of Biology and Biotechnology, Worcester Polytechnic Institute, Worcester, MA 01605, USA
[2]Department of Molecular Biology and Genetics, Cornell University, Ithaca, NY 14853, USA
[3]Department of Chemistry and Biochemistry, Worcester Polytechnic Institute, Worcester, MA 01605, USA
[4]Division of Biology and Biological Engineering, California Institute of Technology, Pasadena, CA 91125, USA
[5]Present address: Merck and Co, 2000 Galloping Hill Road, Kenilworth, NJ 07033, USA
[6]Present address: Individualized Cell and Gene Therapies, South San Francisco, Genentech, CA 94080, USA
[7]Present address: Biology Department, Tufts University, Medford, MA 02155, USA
[8]Bioinformatics and Computational Biology Program, Worcester Polytechnic Institute, Worcester, MA 01605, USA

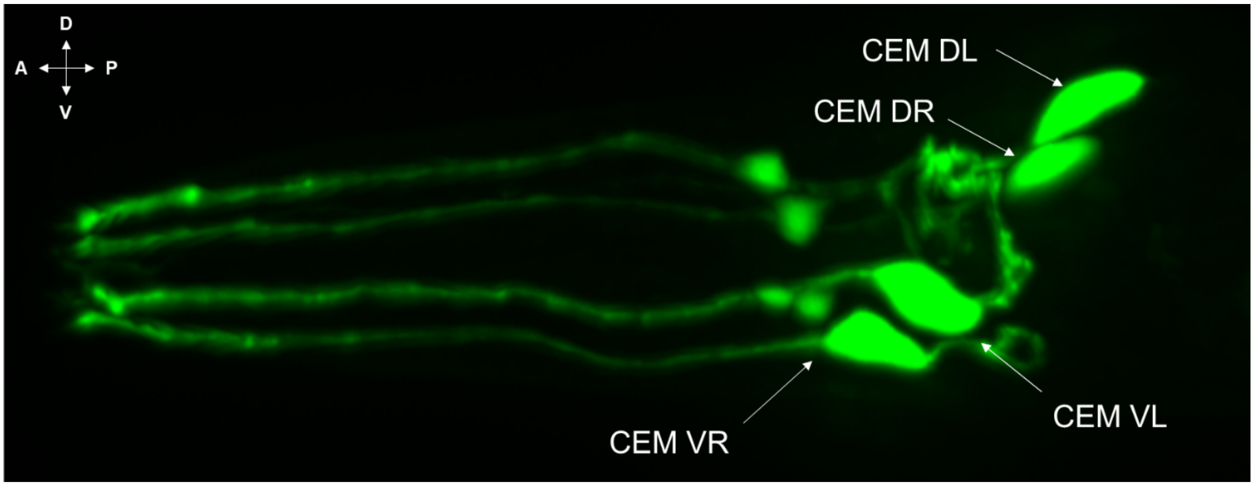


**Figure S1. Expression profile of p*trf-1*::GFP localizes to the CEM neurons.** GFP driven by the promoter of *trf-1* gene localizes in all subcellular compartments of the CEM neurons, including soma, dendrites, and cilia, providing a new avenue for targeting of the male-specific neurons.

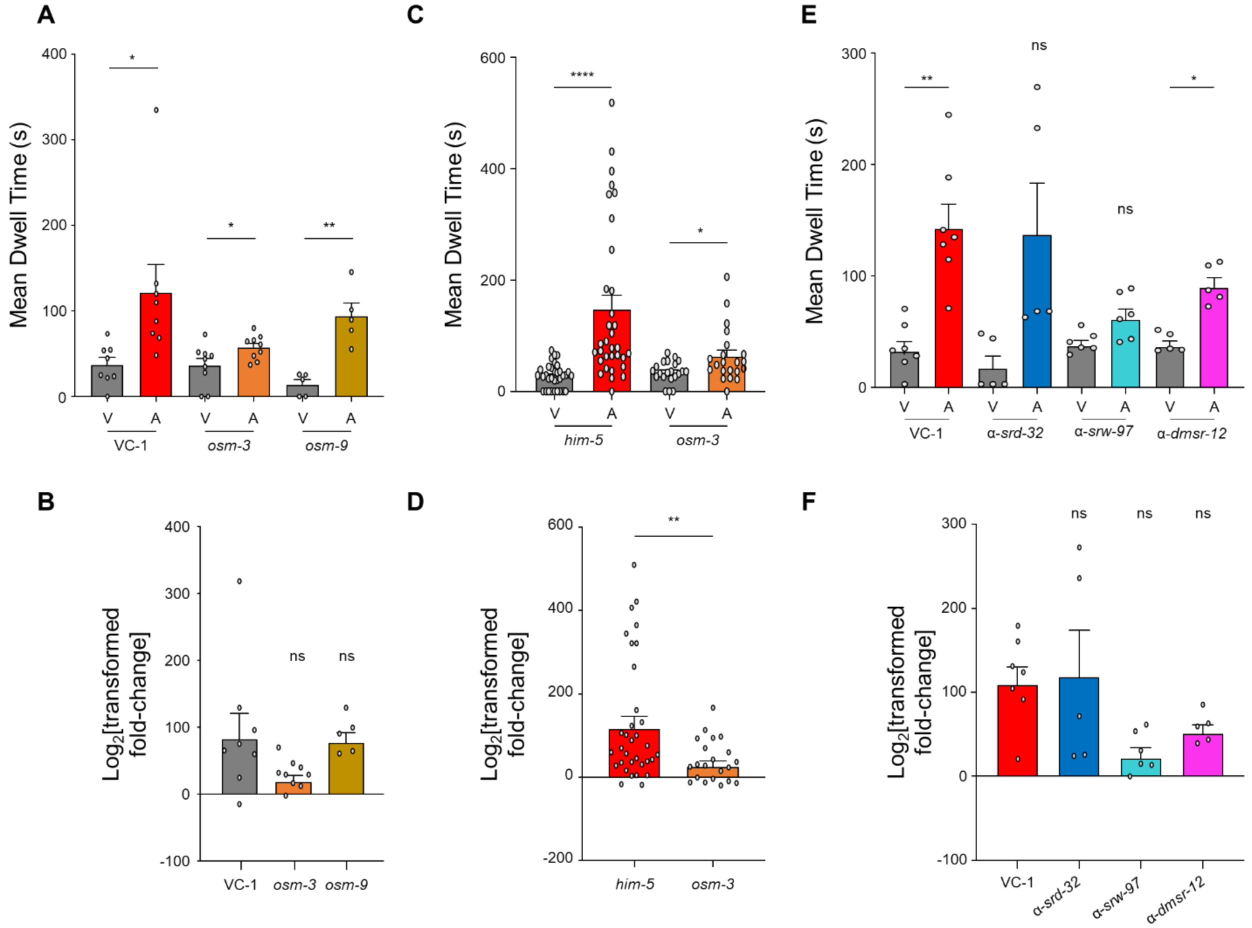


**Figure S2. Knockdown of CEM-enriched GPCR candidates results in aberrant behavioral response to ascr#8. (A)** Raw dwell times and **(B)** Log2(fold-change of ascr#8/vehicle) of RNAi knockdown of the kinesin motor, *osm-3*, and the TRPV channel, *osm-9* confirm the ability of RNAi feeding to affect pheromone-driven behaviors in a targeted manner. $n \geq 5$. 'V' denotes wells with vehicle negative control in their centers; 'A' denotes wells with ascaroside in their centers. **(C)** Raw dwell times and **(D)** Log2(fold-change) of genetic null *osm-3* mutants reveal an inability to respond to ascr#8. $n \geq 17$. **(E)** Raw dwell time and **(F)** Log2(fold-change) of GPCR of interest (*srd-32*, *dmsr*-12, and *srw-97*) knockdown reveals *dmsr-12* and *srw-97* as contributors to the ascr#8 behavioral response. $n \geq 4$. Error bars denote SEM. **(A, C, E)** Paired *t*-tests and Wilcoxon tests of vehicle control versus ascr#8 dwell time, **(B)** ANOVA, followed by Bonferroni post-hoc tests comparing Log2(fold-change) of RNAi knockdowns to VC-1 control and to each other, **(D)** Mann-Whitney test comparing *him-5* and *osm-3* Log2(fold-change) in response to ascr#8, **(F)** ANOVA, followed by Dunnett's post-hoc test comparing Log2(fold-change) of RNAi knockdowns to VC-1 control. * $p < 0.05$, ** $p < 0.01$, **** $p < 0.0001$.

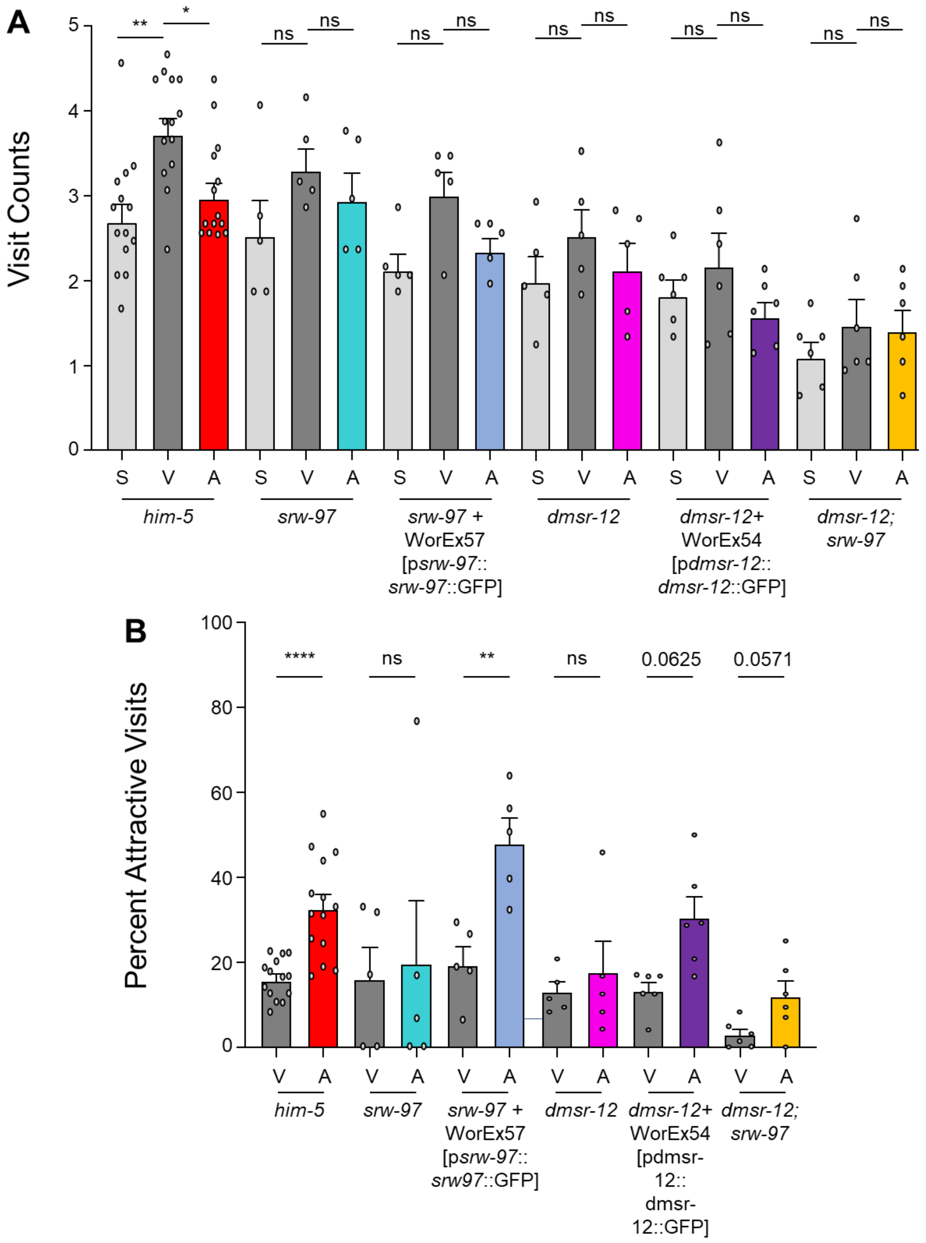


**Figure S3. Effect of genetic knockout and rescue of *srw-97* and *dmsr-12* genes on ascr#8 visit count and attractiveness. (A)** Visit counts of *him-5*, *srw-97* knockout, *srw-97* rescue (WorEx57), *dmsr-12* knockout, *dmsr-12* rescue (WorEx54), and *dmsr-12;srw-97* double knockout males in response to ascr#8. 'S' denotes wells with nothing added to their centers (i.e., a 'spatial' control); 'V' denotes wells with vehicle negative control in their centers; and 'A' denotes wells with ascaroside in their centers. **(B)** Percent of attractive visits per male in the same animals. Light grey denotes spatial controls ("S"), dark grey denotes vehicle controls ("V"), colors denote ascr#8 values ("A") (*him-5*, red; *srw-97*, teal; WorEx57, light blue; *dmsr-12*, pink; WorEx54, purple; *dmsr-12;srw-97*, yellow). Error bars denote SEM**.** $n \geq 5$. **(A)** ANOVA followed by Bonferroni's post-hoc test and Friedman Test followed by Dunn's post-hoc test, **(B)** Paired *t*-test and Wilcoxon matched-pairs signed rank test. * $p < 0.05$, ** $p < 0.01$, *** $p < 0.001$, **** $p < 0.0001$.

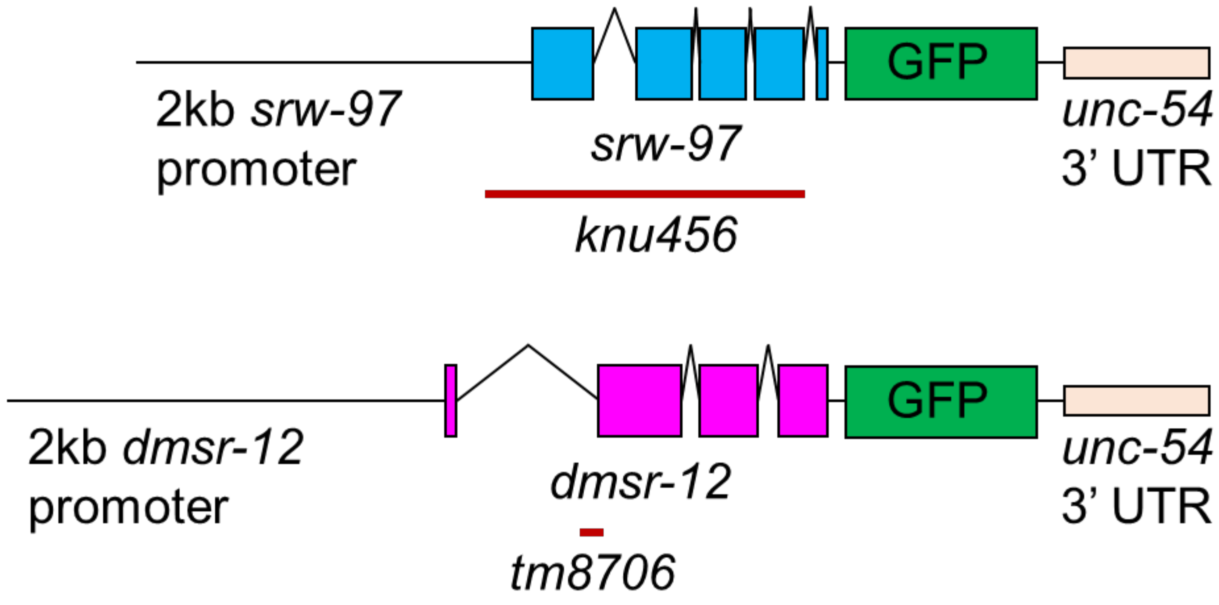


**Figure S4. Generation of *srw-97* and *dmsr-12* rescues.** Genetics maps of *srw-97* (top) and *dmsr-12* (bottom), including the 2 kilobase promoter region used in generation of rescue constructs. The GFP-tag and 3' UTR regions are also displayed. The red lines indicate the 1620 and 118 nucleotide deletion regions of *srw-97 (knu456)* and *dmsr-12 (tm8706)*, respectively.

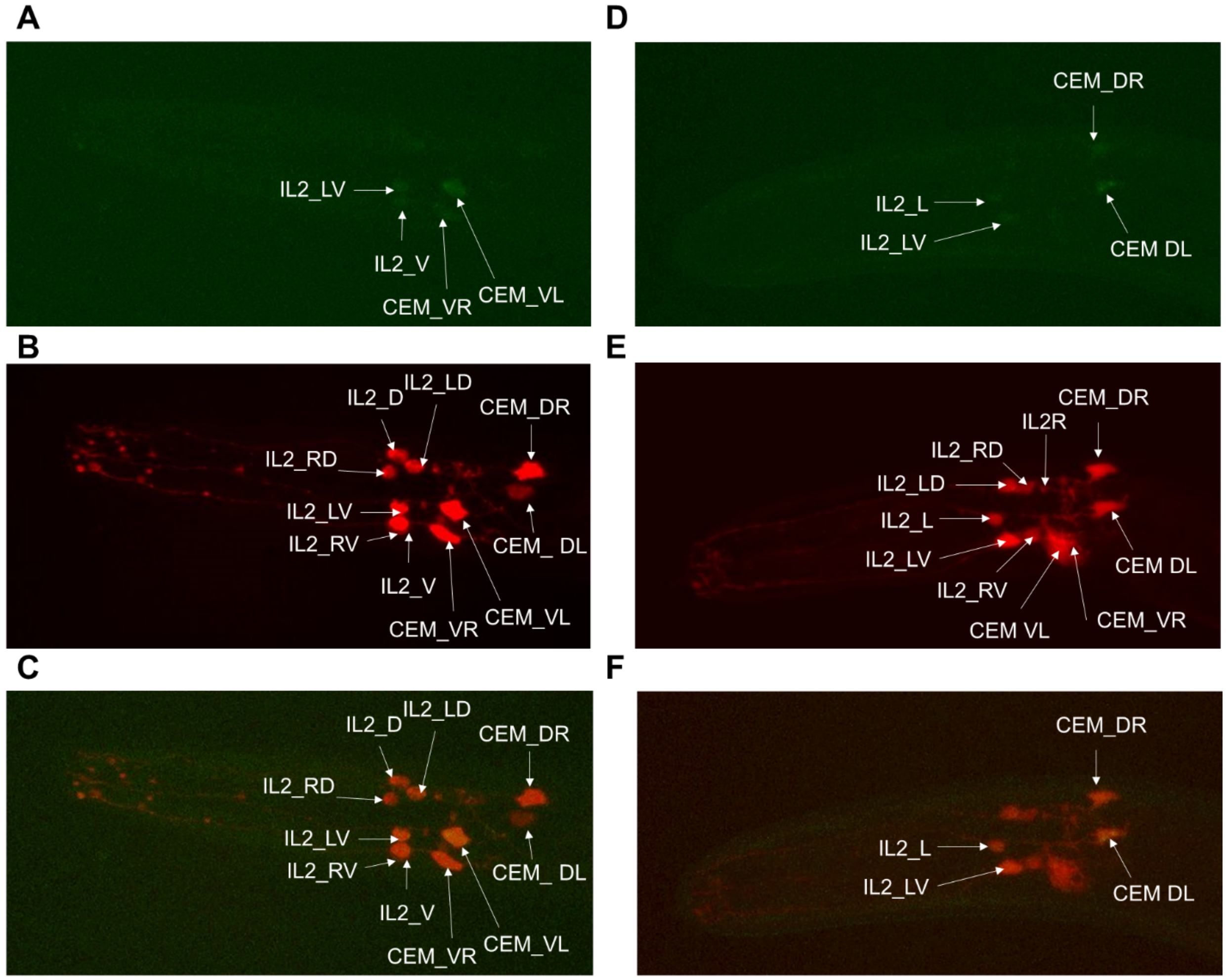


**Figure S5. Expression of *srw-97* and *dmsr-12* within CEM neurons. (A-C)** Expression of WorEx61 (p*srw-97*::*srw-97*::GFP) in myIs20 (p*klp-6*::tdTomato), which expresses in EVN neurons **(B,E)**. The receptor-fusion is observed in CEM_VR, CEM_VL, and IL_V **(A, C)**. **(D-F)** Expression of WorEx60 (p*dmsr-12*::*dmsr-12*::GFP) in myIs20. The receptor-fusion is observed in CEM_DR, CEM_DL, and IL2_L. Both receptors show expression within IL_LV **(A, C, D, F)**. Images captured at 63X using an oil lens.

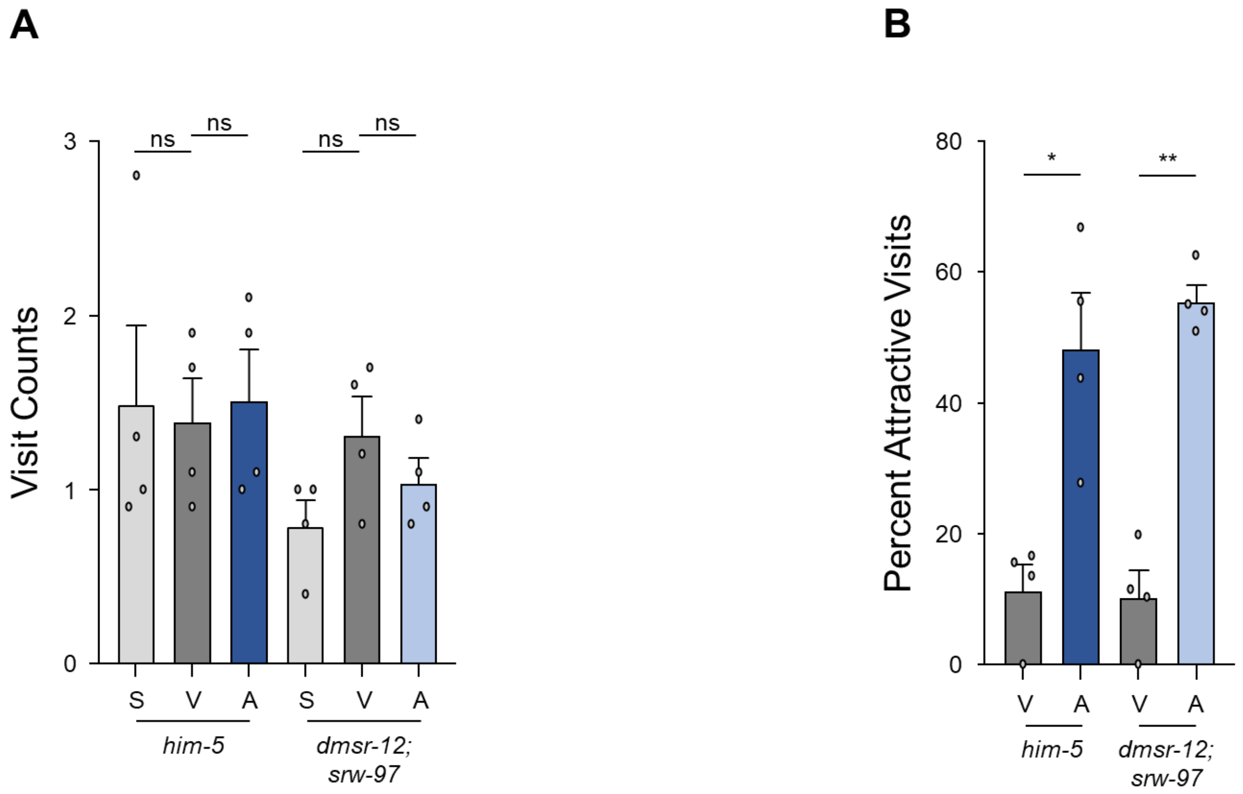


**Figure S6. Loss of both the *dmsr-12* and *srw-97* GPCRs has no effect on the behavioral response to ascr#3. (A)** Visit counts of *him-5* and *dmsr-12;srw-97* double knockout males in response to ascr#3. **(B)** Percent of attractive visits per male in the same animals. Light grey denotes spatial controls ("S"), dark grey denotes vehicle controls ("V"), colors denote ascr#8 values ("A") (*him-5*, dark blue; *dmsr-12*, light blue). Error bars denote SEM**.** $n \geq 5$. **(A)** ANOVA followed by Bonferroni's post-hoc test, **(B)** Paired $t$-test. * $p < 0.05$, ** $p < 0.01$, *** $p < 0.001$, **** $p < 0.0001$.

**Table S1. CEM neuron-specific protein-coding gene counts.**

Protein-coding genes specific to CEMs generally are defined here by being expressed at least two times more strongly in the most strongly expressing CEM type than in the most strongly expressing of three replicates of adult *C. elegans* males. Protein-coding genes enriched in or specific to a particular CEM neuron type are defined here by two traits. To be enriched, they must be expressed at least two times more strongly in a given CEM cell type than the other three CEM cell types (i.e., they must have ≥2-fold higher expression in one CEM type than any of the other three). To be not merely enriched but also specific, they must in addition be expressed at least two times more strongly in a given CEM neuron type than in the most strongly-expressing replicate from three replicates of whole adult *C. elegans* males (i.e., they must have ≥2-fold higher expression in one CEM type than any of three adult *C. elegans* male replicates). Note that it is possible for a gene to be CEM-specific (expressed two-fold more in CEMs than whole adult males) while not being CEM type-specific (expressed two-fold more in one CEM type than in any other, while also being two-fold more than in adult males). Full expression data are given in **Table S5**.

| **CEM neuron type** | **Enriched genes** | **Specific genes** |
|---|---|---|
| CEM (any type) | Not defined | 267 |
| CEM DL | 1,531 | 62 |
| CEM DR | 770 | 40 |
| CEM VL | 162 | 18 |
| CEM VR | 3,156 | 44 |

**Table S2. Numbers of filtered and mapped reads in RNA-seq data sets.**

"Total reads" denotes the complete set of quality-filtered reads for a given data set that were mapped with RSEM to a *C. elegans* gene index (from WormBase release WS245) to compute gene expression values. With RSEM, a read can be mapped to the gene index either 0 times (i.e., it can fail to map at all); it can map exactly 1 time (i.e., it can map to a unique site in the gene index); or it can map 2+ times. For each RNA-seq data set, the numbers and percentages of reads with each status are given, as is the overall percentage of reads that mapped to the gene index. Whole larval RNA-seq data were originally generated by (SCHWARZ *et al.* 2012). Whole adult *C. elegans* male and female RNA-seq data used here were originally generated by (THOMAS *et al.* 2012). Reads were quality-filtered and mapped as described in Methods.

| **Input data (post-filter)** | **Total reads** | **Aligned 0 times** | **Aligned exactly 1 time** | **Aligned 2+ times** | **Overall alignment rate** |
|---|---|---|---|---|---|
| CEM_DL | 60,200,909 | 31,809,012 (52.84%) | 9,230,910 (15.33%) | 19,160,987 (31.83%) | 47.16% |
| CEM_DR | 63,094,511 | 45,826,303 (72.63%) | 5,967,503 (9.46%) | 11,300,705 (17.91%) | 27.37% |
| CEM_VL | 55,902,297 | 33,428,997 (59.80%) | 6,697,075 (11.98%) | 15,776,225 (28.22%) | 40.20% |
| CEM_VR | 73,412,777 | 60,203,068 (82.01%) | 5,153,127 (7.02%) | 8,056,582 (10.97%) | 17.99% |
| Adult male 1 | 23,534,532 | 1,862,829 (7.92%) | 10,165,410 (43.19%) | 11,506,293 (48.89%) | 92.08% |
| Adult male 2 | 27,405,842 | 2492999 (9.10%) | 11,178,629 (40.79%) | 13,734,214 (50.11%) | 90.90% |
| Adult male 3 | 24,957,082 | 2982445 (11.95%) | 10,078,734 (40.38%) | 11,895,903 (47.67%) | 88.05% |
| Adult female 1 | 26,708,038 | 1038807 (3.89%) | 10,956,545 (41.02%) | 14,712,686 (55.09%) | 96.11% |
| Adult female 2 | 28,711,730 | 2059650 (7.17%) | 11,553,368 (40.24%) | 15,098,712 (52.59%) | 92.83% |
| Adult female 3 | 20,174,660 | 2380109 (11.80%) | 7,643,465 (37.89%) | 10,151,086 (50.32%) | 88.20% |
| Larvae | 23,369,056 | 5513339 (23.59%) | 7,422,262 (31.76%) | 10,433,455 (44.65%) | 76.41% |

**Table S3. Numbers of genes with above-background expression in CEM neuronal and larval RNA-seq data.**

Protein-coding and ncRNA-coding genes were counted as detectably expressed if, in a given RSEM analysis of a given data set, the gene had an expression value of ≥ 0.1 transcripts per million (i.e., ≥ 0.1 TPM); they were counted as robustly expressed if the gene had an expression value of ≥ 0.1 TPM in a 99% credibility interval (i.e., ≥ 0.1 minTPM). All gene annotations (including those used here, such as coding status) are given in **Table S5**.

| Data set | Protein-coding, detectably expressed | Protein-coding, robustly expressed | ncRNA-coding, detectably expressed | ncRNA-coding, robustly expressed |
|---|---|---|---|---|
| CEM_DL | 4,728 | 2,179 | 8,894 | 4 |
| CEM_DR | 5,333 | 1,500 | 9,087 | 9 |
| CEM_VL | 3,463 | 935 | 8,995 | 4 |
| CEM_VR | 7,698 | 1,091 | 9,163 | 22 |
| Adult male 1 | 18,263 | 15,486 | 9,119 | 10 |
| Adult male 2 | 18,182 | 15,508 | 9,093 | 18 |
| Adult male 3 | 18,524 | 15,616 | 9,126 | 16 |
| Adult female 1 | 17,699 | 13,440 | 9,147 | 8 |
| Adult female 2 | 17,939 | 14,438 | 9,140 | 9 |
| Adult female 3 | 18,246 | 13,398 | 9,178 | 11 |
| Larvae | 15,023 | 9,922 | 9,151 | 7 |

**Table S4. Proteomes used for phylogenetic analysis of select CEM-expressed genes.**

A phylogeny of the *Caenorhabditis* genus that includes these species and that defines the Elegans and Japonica groups within *Caenorhabditis* has been published (STEVENS *et al.* 2020), updating previous phylogenetic analyses. Genome assemblies for *C. remanei*, *C. brenneri*, and *C. japonica* were generated by the Washington University Genome Center (WashU) and made publicly available by 2010 (HARRIS *et al.* 2010). Other genomes have been published as noted.

| Organism | Features | URL | Reference |
|---|---|---|---|
| *C. elegans* | Hermaphrodite | [For almost all of the proteome:] *ftp://ftp.wormbase.org/pub/wormbase/releases/WS275/species/c_elegans/PRJNA13758/c_elegans.PRJNA13758.WS275.protein.fa.gz*<br><br>[For an earlier long isoform of *dmsr-12*:] *ftp://ftp.wormbase.org/pub/wormbase/releases/WS250/species/c_elegans/PRJNA13758/c_elegans.PRJNA13758.WS250.protein.fa.gz* | (CONSORTIUM 1998) |
| *C. inopinata* | Male-female sibling species of *C. elegans* | *ftp://ftp.wormbase.org/pub/wormbase/releases/WS275/species/c_inopinata/PRJDB5687/c_inopinata.PRJDB5687.WS275.protein.fa.gz* | (KANZAKI *et al.* 2018) |
| *C. briggsae* | Hermaphrodite | *ftp://ftp.wormbase.org/pub/wormbase/releases/WS275/species/c_briggsae/PRJNA10731/c_briggsae.PRJNA10731.WS275.protein.fa.gz* | (STEIN *et al.* 2003) |
| *C. nigoni* | Male-female sibling species of *C. briggsae* | *ftp://ftp.wormbase.org/pub/wormbase/releases/WS275/species/c_nigoni/PRJNA384657/c_nigoni.PRJNA384657.WS275.protein.fa.gz* | (YIN *et al.* 2018) |
| *C. tropicalis* | Hermaphrodite | n/a | Unpublished |
| *C. wallacei* | Male-female sibling species of *C. tropicalis* | n/a | Unpublished |
| *C. remanei* | Male-female, Elegans group | *ftp://ftp.wormbase.org/pub/wormbase/releases/WS275/species/c_remanei/PRJNA53967/c_remanei.PRJNA53967.WS275.protein.fa.gz* | (HARRIS *et al.* 2010) |
| *C. brenneri* | Male-female, Elegans group | *ftp://ftp.wormbase.org/pub/wormbase/releases/WS275/species/c_brenneri/PRJNA20035/c_brenneri.PRJNA20035.WS275.protein.fa.gz* | (HARRIS *et al.* 2010) |
| *C. japonica* | Male-female, Japonica group | *ftp://ftp.wormbase.org/pub/wormbase/releases/WS275/species/c_japonica/PRJNA12591/c_japonica.PRJNA12591.WS275.protein.fa.gz* | (HARRIS *et al.* 2010) |
| *C. becei* | Male-female, Japonica group | *http://download.caenorhabditis.org/v1/sequence/Caenorhabditis_sp29_QG2083_v1.proteins.fa.gz* | (STEVENS *et al.* 2019) |
| *C. sulstoni* | Male-female, Japonica group | *http://download.caenorhabditis.org/v1/sequence/Caenorhabditis_sp32_JU2788_v1.proteins.fa.gz* | (STEVENS *et al.* 2019) |

**Table S5. Traits of *C. elegans* genes expressed in CEM neurons (CEM_DL, CEM_DR, CEM_VL, and CEM_VR), adult males, adult females, and whole larvae.**

Several subtables (sheets) are provided here. **Table S5a**: 'CEM_RNAseq_annots' provides all data for all genes (both protein-coding and ncRNA-coding). **Table S5b**: 'Protein-coding' provides data for all protein-coding genes. 'CEM-specific' provides data for all protein-coding genes specifically expressed in CEMs generally (total numbers, and definition of 'specific', given in **Table S1**). 'CEM DL-enriched', 'CEM DR-enriched', 'CEM VL-enriched', and 'CEM VR-enriched' provide data for all protein-coding genes enriched in each of the four CEM neuron types (total numbers, and definition of 'enriched', given in **Table S1**). 'CEM DL-specific', 'CEM DR-specific', 'CEM VL-specific', and 'CEM VR-specific' provide data for all protein-coding genes specific to each of the four CEM neuron types (total numbers, and definition of 'specific', given in **Table S1**). Finally, **Table S5c**: 'Pseudozeroed TPMs' provides gene expression data with empirical non-zero pseudominimum values (pseudozeroes, denoted with TPM_nz), which allows comparisons without division by zero.

The data columns in each subtable are as follows:

**Gene:** a given predicted protein-coding or ncRNA-coding gene in the *C. elegans* genome, from WormBase release WS245, for which we observed non-zero gene activity in CEM neurons. All further data columns are pertinent to that particular gene.

**[*RNA-seq read set 1*]/[*RNA-seq read set 2*]:** the ratio of gene expression (measured in TPM) between two RNA-seq data sets. In some of these comparisons, 'max' denotes that the maximum gene expression value seen for a set of related data sets being is used. For instance, 'CEM_all.max_TPM/male.max_TPM' compares, for each gene, its highest expression value seen in any CEM neuron type versus its highest expression value seen in any of three biological replicates of adult males; 'CEM_DL_TPM/CEM_non_DL.max_TPM' compares, for each gene, its expression value in a particular CEM type (in this case, DL) versus its highest expression level seen in any of the other three CEM types (in this case, DR, VL, and VR); 'CEM_any/CEM_non_any.max' compares, for each gene, its highest expression value in *any* particular CEM type versus its highest expression level seen in any of the other three CEM types. All expression values are in transcripts per million (TPM) and are provided in [RNA-seq read set]_TPM columns (described further below).

For computing expression ratios, zero gene expression zero values in a given RNA-seq data set were replaced with empirical non-zero pseudominimum values, because replacing zero values with empirical non-zero pseudominima allows logarithmic plotting and comparison of expression ratios for all genes. For a given RNA-seq data set, we defined its empirical non-zero pseudominimum as being the smallest non-zero expression value observed in our RNA-seq analysis, reasoning that the smallest such non-zero value is effectively equivalent to noise.

**Coding:** the nature of a given gene's coding potential, as annotated in WormBase WS245. Most genes are either solely protein-coding or solely ncRNA-coding and are noted as such in this data column. For 301 genes in *C. elegans*, WS245 predicts both protein-coding and non-protein-coding transcripts; in this table, such genes are denoted with "protein; ncRNA".

However, for purposes of gene analysis, we assume that any gene with dual predicted nature is solely protein-coding.

**Prot_size:** the full range of sizes for all protein products from a gene's predicted isoforms.

**Max_prot_size:** the size of the largest predicted protein product.

**Housekeeping:** a set of genes that were previously observed, by single-cell RNA-seq, to be constitutively active both in whole *C. elegans* larvae and in three different developmental stages/genotypes of migrating *C. elegans* linker cells (SCHWARZ *et al.* 2012).

**TF:** genes annotated as encoding transcription factors by one or more of three different censuses by Gupta, Thomas, or Walhout, as previously compiled (SCHWARZ *et al.* 2012).

**7TM_GPCRs:** a set of genes encoding G-protein coupled receptors (GPCRs), a class of genes of particular biological interest in deciphering CEM function (HOBERT 2013).

**PFAM:** for protein-coding genes, predicted domains from Pfam 31.0 (FINN *et al.* 2016), with a domain-specific calibrated significance threshold (invoked by the *hmmscan* argument *--cut_ga*).

**GO_term:** Gene Ontology terms (GENE ONTOLOGY 2015) for which a gene was annotated in WormBase release WS245.

**eggNOG:** for protein-coding genes, predicted orthology groups from the eggNOG 3.0 database (POWELL *et al.* 2012).

**Phobius:** predictions of signal and transmembrane sequences made with Phobius (KÄLL *et al.* 2004). 'SigP' indicates a predicted signal sequence, and 'TM' indicates one or more transmembrane-spanning helices, with N helices indicated with '(Nx)'. Varying predictions from different isoforms are listed.

**NCoils:** coiled-coil domains, predicted by ncoils (LUPAS 1996). As with Psegs, the relative and absolute fractions of each protein's coiled-coil residues are shown.

**Psegs:** this shows what fraction of a protein is low-complexity sequence, as detected by pseg (WOOTTON 1994). Both the proportion of such sequence (ranging from 0.01 to 1.00) and the exact ratio of low-complexity residues to total residues are given. Proteins with no predicted low-complexity residues are blank.

**[*RNA-seq read set*]_TPM:** for a given RNA-seq data set, this denotes the expression level in transcripts per million (TPM) for a given gene. The read sets include particular CEM neuron types (CEM_DL, CEM_DR, CEM_VL, or CEM_VR; **Table S2**), whole larvae (generated from a pooled set of all larval RNA-seq reads; **Table S2**), three biological replicates of adult *C. elegans* males (**Table S2**), and three biological replicates of adult *C. elegans* females (**Table S2**).

**[*RNA-seq read set*]_minTPM:** for the gene in question, and for a given RNA-seq data set, this denotes the minimum estimate of that gene's activity as measured in that RNA-seq data set and computed by RSEM with a 99% confidence interval in transcripts per million (minTPM). All cell types are as with "[*RNA-seq read set*]_TPM" above.

**[*RNA-seq read set*]_[optional modifier]TPM_nz:** for the gene in question, and for a given RNA-seq data set, this denotes for a given RNA-seq data set from either a particular CEM neuron type or from larvae, this denotes the same expression level for a given gene as in [RNA-seq read set]_TPM above, but with any zero gene expression values in a given RNA-seq data set replaced with empirical non-zero pseudominimum values. See '[RNA-seq read set 1]/[RNA-seq read set 2]' above for further explanations of this point. All cell types are as with "[*RNA-seq read set*]_TPM" above.

**[*RNA-seq read set*]_reads:** for the gene in question, and for a given RNA-seq data set, this denotes a posterior mean estimate of the number of RNA-seq reads mapping to that gene as computed by RSEM, and with decimal fractions rounded off. All cell types are as with "[*RNA-seq read set*]_TPM" above.

**Table S6. Coding Sequences Included in GPCR-GFP Fusion Constructs.**
The percentage of codons from each GPCR gene coding sequence included in the GFP transgenes varied from construct to construct, although the majority was included in each product.

| Gene (Transcript) | Promoter Length (nt) | Percentage of Codons Included in GFP Fusion (included/total amino acids) |
|---|---|---|
| *seb-3* (C18B12.2.1) | 3,082 | 100% (454/454) |
| *dmsr-12* (H34P18.1a) | 2,810 | 99.1% (360/363) |
| *srw-97* (ZC204.15) | 1,360 | 99.7% (371/372) |
| *srd-32* (T19H12.5) | 1,874 | 52.8% (179/339) |
| *srr-7* (T01G5.4) | 657 | 69.5% (269/387) |
| *trf-1* (F45G2.6.1) | 4,044 | N/A |

**Table S7. Quantification of SRW-97::GFP and DMSR-12::GFP in Rescue Strains.**

Average fluorescence (arbitrary units, background subtracted) of SRW-97::GFP and DMSR-12::GFP calculated by cell. Values displayed as Average (SEM). n > 10.

| | CEM_DL | CEM_DR | CEM_VL | CEM_VR |
|---|---|---|---|---|
| DMSR-12::GFP | **11.14** (2.99) | 4.55 (1.19) | 2.30 (1.76) | 1.84 (0.76) |
| SRW-97::GFP | 6.07 (1.75) | 4.95 (1.43) | **10.44** (3.02) | 6.81 (1.97) |

**Table S8. Strains**
Strains utilized in this study.

| Figure | Strain | Genotype | Source |
|---|---|---|---|
| - | CU607 | smIs23 [p*pkd-2*::GFP+ pBX]; *him-5(e1490)* | Ding Xue |
| 2A | - | bsIs14 [p*pkd-2::*GFP + *pha-1*(+)]*;him-5(e1490)* | Doug Portman |
| - | - | *pha-1(e2123ts);lite-1(ce314);him-5(e1490)* | Rene Garcia |
| 2B | JSR40 | *pha-1(e2123ts);lite-1(ce314);him-5(e1490);* worEx8 [p*seb-3::*GFP; *pha-1*(+)] | InVivo Biosystems/ DKR |
| 2C | JSR36 | *pha-1(e2123ts);lite-1(ce314);him-5(e1490);* worEx4 [p*dmsr-12::*GFP; *pha-1*(+)] | InVivo Biosystems/ DKR |
| 2D | JSR13 | *pha-1(e2123ts);lite-1(ce314);him-5(e1490);* worEx02 [p*srd-32::*GFP; *pha-1*(+)] | InVivo Biosystems/ DKR |
| 2E | JSR15 | *pha-1(e2123ts);lite-1(ce314);him-5(e1490);* worEx03 [*psrw-97::*GFP; *pha-1*(+)] | InVivo Biosystems/ DKR |
| 2F | JSR10 | *pha-1(e2123ts);lite-1(ce314);him-5(e1490);* worEx01 [p*srr-7::*GFP; *pha-1*(+)] | InVivo Biosystems/ DKR |
| Fig. S1 | JSR39 | *pha-1(e2123ts);lite-1(ce314);him-5(e1490);* worEx07 [p*trf-1::*GFP; *pha-1*(+)] | InVivo Biosystems/ DKR |
| - | VH624 | rhIs13 [*unc-119::*GFP + dpy-20(+)]*;nre-1(hd20);lin-15B(hd126)* | *Caenorhabditis* Genetics Center (CGC) via Erich Schwarz |
| Fig. S2 | JSR44 | rhIs13 [*unc-119::*GFP + dpy-20(+)]*;nre-1(hd20);lin-15B(hd126);him-5(e1490)* | DKR |
| Fig 3, Fig. S2, Fig S3, Fig. S6 | CB4088 | *him-5(e1490)* | CGC |
| Fig 3, Fig. S3 | JSR55 | *srw-97(knu456);him-5(e1490)* | InVivo Biosystems/ DKR |
| Fig 3, Fig. S3, Fig. S5 | JSR158 | *srw-97(knu456);him-5(e1490);* WorEx57[p*srw-97::srw-* | InVivo Biosystems/ CSM |

| | | | |
|---|---|---|---|
| | | *97::*GFP*:: unc54-3'* UTR*; unc-122::*RFP*]* | |
| Fig 3, Fig. S5 | JSR179 | myIs20 [p*klp-6*::*tdTomato* + pBx]; *pha-1(e2123)*; *him-5(e1490)*; WoEx61[*srw-97*::GFP::*unc54*-3' UTR; *unc-122*::RFP] | CSM |
| - | tm8706 | *dmsr-12(tm8706)* | National BioResource Project |
| Fig 3, Fig. S3 | JSR68 | *dmsr-12(tm8706);him-5(e1490)* | DKR |
| Fig 3, Fig. S3 | JSR149 | *dmsr-12(tm8706);him-5(e1490);* WorEx54[p*dmsr-12::dmsr-12::*GFP*:: unc54-3'* UTR*; unc-122::*RFP*]* | InVivo Biosystems/ CSM |
| Fig 3, Fig. S5 | JSR178 | myIs20 [p*klp-6*::*tdTomato* + pBx]; *pha-1(e2123)*; *him-5(e1490)*; WorEx60[*dmsr-12*::GFP::*unc54*-3' UTR; *unc-122*::RFP] | CSM |
| Fig 3, Fig. S3, Fig. S6 | JSR70 | *srw-97(knu456);dmsr-12(tm8706);him-5(e1490)* | ANM |

**Table S9. Primers**
Primers used to generate GFP-promoter fusions in **Figure 2** and **Fig. S1**, and rescue strains in **Figure 3** and **Fig. S3-S5**.

| Figure | Primer | |
|---|---|---|
| | Name | Sequence |
| Fig. 2B | seb-3_A | TTGACAGTAACTGGCGCTAC |
| Fig. 2B | seb-3_B | AGTCGACCTGCAGGCATGCAAGAGATTTCGTAGACACCGA GTAGAT |
| Fig. 2B | seb-3_Ap | ACAGTAACTGGCGCTACTCCC |
| Fig. 2C | dmsr-12_A | ATTTCCCCAAGGAGTTTTCA |
| Fig. 2C | dmsr-12_B | AGTCGACCTGCAGGCATGCAAGGAAGCGTGTGCAACTGAA TG |
| Fig. 2C | dmsr-12_Ap | CCCAAGGAGTTTTCAATCTT |
| Fig. 2D | srd-32_A | TTTTTGTGGATTTTGTTCCA |
| Fig. 2D | srd-32_B | AGTCGACCTGCAGGCATGCAAGCTGGTTCCCACCACAGTCC TATA |
| Fig. 2D | srd-32_Ap | TGTGGATTTTGTTCCAATGA |
| Fig. 2E | srw-97_A | GTCGGAAAACTCAAAAAGCA |
| Fig. 2E | srw-97_B | AGTCGACCTGCAGGCATGCAAGCTTCGATTTGACGTGCTGC TGTG |
| Fig. 2E | srw-97_Ap | GGAAAACTCAAAAAGCAAACA |
| Fig. 2F | srr-7_A | ACATTTGGAAAGCGTAGAAAA |
| Fig. 2F | srr-7_B | AGTCGACCTGCAGGCATGCAAGCTGAAGTCAAGTAGACGC CGTTT |
| Fig. 2F | srr-7_Ap | TTGGAAAGCGTAGAAAATTGA |
| Fig. 2B-F, Fig. S1 | GFP_C | AGCTTGCATGCCTGCAGGTCGACT |
| Fig. 2B-F, Fig. S1 | GFP_D | AAGGCCCCGTACGGCCGACTAGTAGG |
| Fig. 2B-F, Fig. S1 | GFP_Dp | GGAAACAGTTATGTTTGGTATATTGGG |
| Fig. S1 | trf-1_A | ATGGCTGACAAGTTGTTCTCG |
| Fig. S1 | trf-1_B | AGTCGACCTGCAGGCATGCAAGTTTCTATGGAATTCAGAAA TTGCT |
| Fig. S1 | trf-1_Ap | GGAGAGGCTTTGGTGAGAAA |
| Fig. 3, Fig. S3-S6 | srw-97 _fwd | TGAAATAAGCTTGCATGCCTGCAGATCGTCATCGGGATCTT C |

| | | |
|---|---|---|
| Fig. 3, Fig. S3-S6 | srw-97_rev | TCCTTTGGCCAATCCCGGGGATCCTACTCGATTTGACGTGCTG |
| Fig. 3, Fig. S3-S6 | GFP_srw_fwd | CACAGCAGCACGTCAAATCGAGTAGGATCCCCGGGATTGGCC |
| Fig. 3, Fig. S3-S6 | GFP_srw_rev | AGAGGTGAAGATCCCGATGACGATCTGCAGGCATGCAAGCTTATTTC |
| Fig. 3, Fig. S3-S6 | dmsr_fwd | ATGAAATAAGCTTGCATGCCTGCAGCCTGAAACGTTGAATGTATTCTAGG |
| Fig. 3, Fig. S3-S6 | dmsr_rev | GTCCTTTGGCCAATCCCGGGGATCCAAATGGTCGCGAAGCGTG |
| Fig. 3, Fig. S3-S6 | GFP_dmsr_fwd | AGTTGCACACGCTTCGCGACCATTTGGATCCCCGGGATTGGCC |
| Fig. 3, Fig. S3-S6 | GFP_dmsr_rev | CCTAGAATACATTCAACGTTTCAGGCTGCAGGCATGCAAGCTTATTTC |

### Supplemental Information References

Consortium, T. *C. e.* S., 1998 Genome sequence of the nematode C. elegans: a platform for investigating biology. Science 282**:** 2012-2018.

Finn, R. D., P. Coggill, R. Y. Eberhardt, S. R. Eddy, J. Mistry *et al.*, 2016 The Pfam protein families database: towards a more sustainable future. Nucleic acids research 44**:** D279-D285.

Gene Ontology, C., 2015 Gene Ontology Consortium: going forward. Nucleic acids research 43**:** D1049-D1056.

Harris, T. W., I. Antoshechkin, T. Bieri, D. Blasiar, J. Chan *et al.*, 2010 WormBase: a comprehensive resource for nematode research. Nucleic acids research 38**:** D463-D467.

Hobert, O., 2013 The neuronal genome of *Caenorhabditis elegans*. WormBook**:** 1-106.

Käll, L., A. Krogh and E. L. L. Sonnhammer, 2004 A combined transmembrane topology and signal peptide prediction method. Journal of molecular biology 338**:** 1027-1036.

Kanzaki, N., I. J. Tsai, R. Tanaka, V. L. Hunt, D. Liu *et al.*, 2018 Biology and genome of a newly discovered sibling species of *Caenorhabditis elegans*. Nature communications 9**:** 3216-3216.

Lupas, A., 1996 Prediction and analysis of coiled-coil structures. Methods in enzymology 266**:** 513-525.

Powell, S., D. Szklarczyk, K. Trachana, A. Roth, M. Kuhn *et al.*, 2012 eggNOG v3.0: orthologous groups covering 1133 organisms at 41 different taxonomic ranges. Nucleic acids research 40**:** D284-D289.

Schwarz, E. M., M. Kato and P. W. Sternberg, 2012 Functional transcriptomics of a migrating cell in Caenorhabditis elegans. Proceedings of the National Academy of Sciences of the United States of America 109**:** 16246-16251.

Stein, L. D., Z. Bao, D. Blasiar, T. Blumenthal, M. R. Brent *et al.*, 2003 The genome sequence of Caenorhabditis briggsae: a platform for comparative genomics. PLoS Biol 1**:** E45.

Stevens, L., M.-A. Félix, T. Beltran, C. Braendle, C. Caurcel *et al.*, 2019 Comparative genomics of 10 new *Caenorhabditis* species. Evolution Letters 3**:** 217-236.

Stevens, L., S. Rooke, L. C. Falzon, E. M. Machuka, K. Momanyi *et al.*, 2020 The Genome of Caenorhabditis bovis. Current biology : CB**:** S0960-9822(0920)30118-30114.

Thomas, C. G., R. Li, H. E. Smith, G. C. Woodruff, B. Oliver *et al.*, 2012 Simplification and desexualization of gene expression in self-fertile nematodes. Curr Biol 22**:** 2167-2172.

Wootton, J. C., 1994 Non-globular domains in protein sequences: automated segmentation using complexity measures. Computers & chemistry 18**:** 269-285.

Yin, D., E. M. Schwarz, C. G. Thomas, R. L. Felde, I. F. Korf *et al.*, 2018 Rapid genome shrinkage in a self-fertile nematode reveals sperm competition proteins. Science 359**:** 55-61.
